# GRAS Family Transcription Factor Binding Behaviors in Sorghum bicolor, Oyrza, and Maize

**DOI:** 10.1101/2024.09.23.614502

**Authors:** Nicholas Gladman, Sunita Kumari, Audrey Fahey, Michael Regulski, Doreen Ware

**Affiliations:** USDA-ARS Robert Holley Center, Ithaca, NY 14853; Cold Spring Harbor Laboratory, Cold Spring Harbor, NY 11724

**Keywords:** GRAS, Transcription Factor, DAP-seq, Sorghum, Maize, Oryza

## Abstract

Identifying non-coding regions that control gene expression has become an essential aspect of understanding gene regulatory networks that can play a role in crop improvements such as crop manipulation, stress response, and plant evolution. Transcription Factor (TF)-binding approaches can provide additional valuable insights and targets for reverse genetic approaches such as EMS-induced or natural SNP variant screens or CRISPR editing techniques (e.g. promoter bashing). Here, we present the first ever DAP-seq profiles of three GRAS family TFs (SHR, SCL23, and SCL3) in the crop *Sorghum bicolor*, *Oryza sativa japonica*, and *Zea mays*. The binding behaviors of the three GRAS TFs display unique and shared gene targets and categories of previously characterized DNA-binding motifs as well as novel sequences that could potentially be GRAS family-specific recognition motifs. Additional transcriptomic and chromatin accessibility data further facilitates the identification of root-specific GRAS regulatory targets corresponding to previous studies. These results provide unique insights into the GRAS family of TFs and novel regulatory targets for further molecular characterization.

## Introduction

GRAS transcription factors (TFs) form a large family of plant-specific TFs. Named from three members of the family, GIBBERELLIN-ACID INSENSITIVE (<u>G</u>AI), REPRESSOR of GA1 (<u>R</u>G<u>A</u>), and SCARECROW (<u>S</u>CR), these TFs serve a multitude of developmental and environmental response functions and comprise several subfamilies and number in the several dozens of individual proteins across plant lineages(Jaiswal et al., 2022). Members of the GRAS contingent have been shown to be involved in meristem development and axial initiation in tomato, petunia, and arabidopsis(Schumacher et al., 1999; Stuurman et al., 2002; Goldy et al., 2021); influencing meiotic progression in pollen development (Morohashi et al., 2003); fruit development and ripening(Huang et al., 2015; Liu et al., 2021b); seed germination(Lim et al., 2013); arbuscular mycorrhizal symbiosis(Gobbato et al., 2012; Floss et al., 2013; Xue et al., 2015); and light signal transduction as well as plant growth and fertility(Peng et al., 1999; Fukazawa et al., 2014; Fukazawa et al., 2017). This broad onus for GRAS TFs is due, in part, to its expansive cross-talk with numerous hormone signaling pathways, including gibberellic acid(Peng et al., 1997; Silverstone et al., 1998; Dill and Sun, 2001; King et al., 2001; Fu et al., 2002; Niu et al., 2019), jasmonic acid(Hou et al., 2010), brassinosteroids(Tong et al., 2009; Tong et al., 2012), and auxin(Gao et al., 2004; Sánchez et al., 2007); which also relates to GRAS genes having involvement in numerous stress responses like drought, heat, salinity, cold(Ma et al., 2010; Yang et al., 2011; Yuan et al., 2016), light(Chen et al., 2015), disease resistance(Fode et al., 2008; Wild et al., 2012; Li et al., 2018), and flavonoid production(Pillet et al., 2015; Huang et al., 2021). Initially, it was shown that many GRAS TFs might require the interaction of other proteins like Indeterminate Domain (IDD) TFs to regulate transcription(Welch et al., 2007; Hirano et al., 2017; Aoyanagi et al., 2020), but other structural studies demonstrated the innate ability of certain GRAS TFs to bind DNA without heterodimerization(Li et al., 2016).

Gene regulatory networks (GRNs) have been useful to identify modules that influence plant growth and development(Tu et al., 2020; Zhu et al., 2023; Fu et al., 2024; Khan et al., 2024). Incorporating multiple -omics datasets into these networks improves the power and resolution of their conclusions. While gene expression, epigenetic, and phenotypic profiling information have been useful to unveil regulatory schema, the addition of TF binding information and DNA-binding motif fingerprinting can add significant improvements to GRN construction and interpretation(Savadel et al., 2021; Shojaee and Huang, 2023) and bioengineering targets(Rodríguez-Leal et al., 2017; Liu et al., 2021a; Cao et al., 2022; Yang et al., 2023). Despite the known importance that many GRAS TFs play throughout plants, few TF binding experiments have been conducted on members of this family in model or crop species(Yoon et al., 2016; Tu et al., 2020). By generating TF binding profiles of GRAS proteins, selected regulatory candidate promoters, enhancers, and gene targets can be identified and modified via CRISPR editing for breeding efforts pertaining to root and shoot development(Ron et al., 2014; Triozzi et al., 2021). These targeted approaches can show greater phenotypic variability than creating null or hypomorphic alleles of the TFs themselves(Aguirre et al., 2023).

Sorghum [*Sorghum bicolor (L.) Moench*] is a globally important C_4_ grass crop with observed drought, heat, and high-salt tolerances with a completely sequenced genome (2x=2n=10; ∼720 Mb)(Paterson et al., 2009; McCormick et al., 2018; Cooper et al., 2019). There are significant discrepancies in understanding the targets and regulatory regions of important TF families in this monocot despite the wealth of genetic resources such as natural diversity panels(Casa et al., 2008) and EMS-mutagenized populations(Jiao et al., 2016; Addo-Quaye et al., 2017) to be used for forward genetics and functional characterization(Jiao et al., 2017; Jiao et al., 2018; Dampanaboina et al., 2019; Gladman et al., 2019) as well as increased genomic profiling of sorghum root, leaf, flower, and seed. In the work presented here, we demonstrate the first DAP-seq profiles of three GRAS family TFs: SHORT ROOT (SHR), SCARECROW-LIKE23 (SCL23), and SCARECROW-LIKE3 (SCL3) in *Sorghum bicolor*; characterize their binding behavior in maize (B73) and oryza (Nipponbare); and demonstrate the identification of conserved binding sites in both promoter and intergenic space through the incorporation of publicly available histone methylation data from root tissues. Further, we extend the potentially novel TF binding motifs discovered through the DAP-seq pipeline via a cross-species projection model using position weight matrices (PWM) to improve the validation of new binding sites. Ultimately, this combined information strengthens the model that some GRAS TFs can bind DNA without interacting with other TFs while also strengthening tissue-specific candidates for genome editing approaches and can help further refine the functions of GRAS protein GRNs within the root system.

## Results

### GRAS Family Transcription Factor Selection and Expression

TF profiling was conducted via DNA Affinity Purification (DAP-seq)(O’Malley et al., 2016) using *Sorghum bicolor* BTx623 gDNA. Three GRAS family TFs were chosen to be profiled via DAP-seq based on their ability to be stably expressed and remain soluble in bacterial expression systems and to represent different clades within the GRAS family as defined by Fan et al. 2021: *SHR* (SORBI_3001G327900) from the SHR clade, *SCL23* (SORBI_3002G342800) from the SCR clade, and *SCL3* (SORBI_3005G029600) from the SCL3 clade. Sufficient levels of protein were able to be generated for DNA pulldowns with the addition of mannitol in the bacterial cultures (**Figure 1a**). Bound DNA fragments were eluted, sequenced, and peaks were called using input DNA fragments as controls.

**Figure 1.**
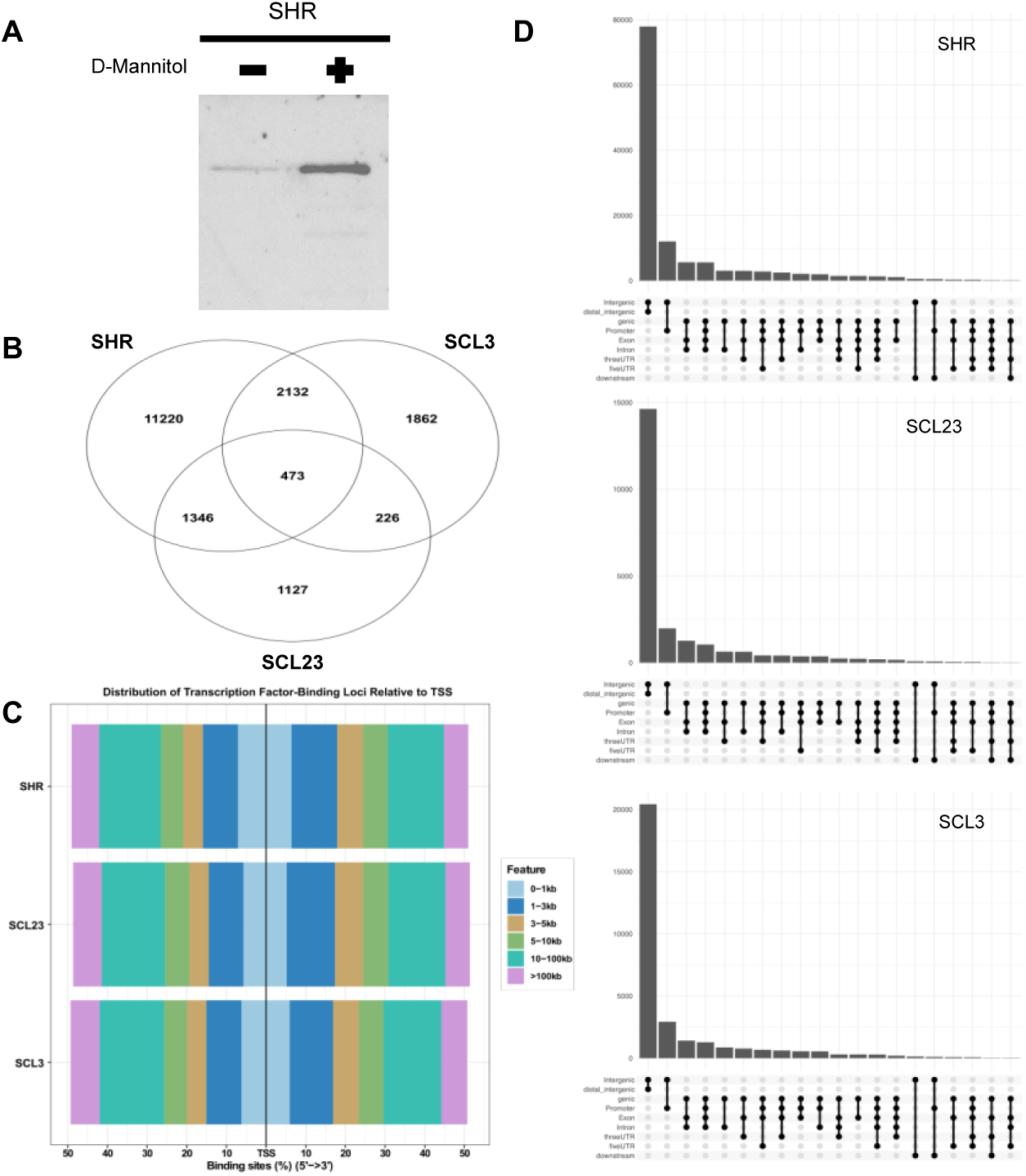
GRAS family DAP-seq results in *Sorghum bicolor.* A) Immunoblot of Sorghum GST-tagged SHR protein eluted of affinity beads. Proteins were induced, isolated, and bound to affinity beads either in the absence or presence (0.10 M) of D-Mannitol. B) All genes with GRAS peaks in their promoter region called from the three DAP-seq pulldowns for Sorghum SHR, SCL23, and SCL3 mapped to the BTx623 v3 genome. C). Distribution of all significant peaks for SHR, SCL23, and SCL3 relative to gene transcriptional start sites. D) Upset plots of the share of genomic features where SHR, SCL23, and SLC3 are binding within the BTx623 genome.

### GRAS Transcription Factor Behaviour in Sorghum bicolor

The peak calling from the SHR, SCL23, and SCL3 DAP-seq yielded tens of thousands of peaks genome-wide for all three TFs, with SHR having the most at >100,000 significant peaks throughout the genome (**Supplemental Data Files 1,2, and 3**). A total of 473 genes had all three GRAS TF binding events in their promoter region (<2,000 bp from transcriptional start site (TSS) to 1000bp after the TSS) (**Figure 1b)**. Most peaks for all three TFs were contained within intergenic space, but around 20% of peaks could be discreetly classified to be within proximal promoter regions (**Figure 1c).** Greater than 6000 peaks between SHR, SCL23, and SCL3 overlapped at least somewhat throughout the genome (**Figure 1d**). While not much DNA-binding information is available for GRAS family proteins in monocots to compare against our results, there are ChIP-chip data from Arabidopsis for SHR, SCL23, and SCL3 orthologs (Cui et al., 2014) that was used for comparison. Based on this, there is ∼60% overlap with the Arabidopsis genes that have peaks within the promoter of a corresponding sorghum ortholog gene and 75% overlap when including distal peaks (2kb-25Kb upstream from the TSS).

When evaluating functional ontologies for genes with GRAS peaks, it was determined that genes with multiple DAP-seq peaks within the promoter did show enrichment in several interesting categories. Namely, gene promoters with multiple SHR peaks in this 3000 bp promoter window had ontologies associated with root hair development (GO:0080147), salicylic acid mediated signaling (GO:2000031), sulfate transport (GO:1902358), hypoxia detection (GO:0070483), amino acid biosynthesis (GO:0000162, GO:0055129), acyl-CoA metabolism (GO:0006637), menaquinone biosynthesis (GO:0009234), and others. The SCL23 cohort of gene promoters with multiple peaks include biological process of electron transport coupled proton transport (GO:0015990), acyl-CoA metabolism (GO:0006637), amino acid and nutrient transport (GO:0015808, GO:1903401, GO:0015813, GO:0009749, GO:0070574, GO:0071805), and very long-chain fatty acid and sphingolipid biosynthesis (GO:0042761, GO:0006665). Similar to SCL23, the genes with multiple SCL3 peaks had ontological processes of electron transport coupled proton transport (GO:0015990) and amino acid transport (GO:0015808) as well as glutathione metabolic process (GO:0006749), cell wall biogenesis (GO:0042546), and cellular oxidant detoxification (GO:0098869). When evaluating all the biological process ontologies of the nearest genes annotated as being associated with a region where all three TFs have overlapping peaks, there is an enrichment for detection of abiotic stimulus (GO:0009582), triterpenoid biosynthetic process (GO:0016104), polyketide biosynthesis (GO:0030639), and phosphorelay signal transduction (GO:0000160). Most of the gene targets accounting for the polyketide biosynthesis ontology are chalcone synthases, which have been shown to be influenced by upstream GRAS activity(Pillet et al., 2015; Huang et al., 2021).

When evaluating whether GRAS peaks exist in important root developmental and stress response pathways, we found that all three TFs had peaks associated with genes involved in gibberellic acid, jasmonic acid, phosphate starvation response, and arbuscular mycorrhizal symbiosis. Notably, either SHR, SCL23, or SCL3 have peaks associated with 89% of all sorghum GRAS TFs; at least one of the three TFs have a peak associated with another GRAS family (**Supplemental Data Table 1**).

### Sorghum GRAS Transcription Factor Binding in Maize and Oryza

To evaluate the consistency of binding targets of the sorghum GRAS TFs in other monocots, we used maize (B73) and oryza (Nipponbare) DNA in the DAP-seq pulldowns using the sorghum SHR, SCL23, and SCL3 proteins. For both Nipponbare and B73, there were fewer peaks called for all three GRAS TFs compared to the sorghum BTx623 gDNA pulldowns (**Figure 2a,b,d,e and Supplemental Data Table 2 and 3**). When focusing on the sorghum genes with GRAS peaks in the promoter, SHR shared 76 gene orthologs with annotated peaks in B73 and 17 in Nipponbare, SCL23 shared 15 gene orthologs with annotated peaks in B73 and 9 in Nipponbare, and SCL3 shared 22 gene orthologs with annotated peaks in B73 and 6 in Nipponbare. This disparity in peak enrichment using the SbGRAS proteins could be due to the divergence in protein sequence identity between the closest orthologs of SbSHR, SbSCL23, and SbSCL3 in oryza and maize: SHR ortholog sequence identity is ∼72-87%; SCL23 ortholog sequence identity is ∼85-97%; and SCL3 ortholog identity is ∼59-67%. Only the SbSCL3 peaks in B73 were enriched for DNA-binding motifs including bHLH, Myb, bZIP and MADS-box. All three sorghum GRAS TF peaks in Nipponbare were enriched for TF-binding motifs. SbSHR had bZIP, LOB, GATA, ABI3, B3, and TCP motifs. SbSCL23 promoter peaks were enriched for WRKY, NAC, GATA, Myb and GRF motifs. SbSCL3 oryza promoter peaks were enriched for ABI3, WRKY, bZIP, Myb and GRF motifs (**Supplemental Data File 4 and 5**). The SHR, SCL23 and SCL3 motif enrichment analyses in either oryza or maize also yielded uncharacterized motifs that shared similar promoter frequency profiles to those identified from the DAP-seq pulldowns that used sorghum gDNA (see following section) **(Figure 2c,f)**.

**Figure 2.**
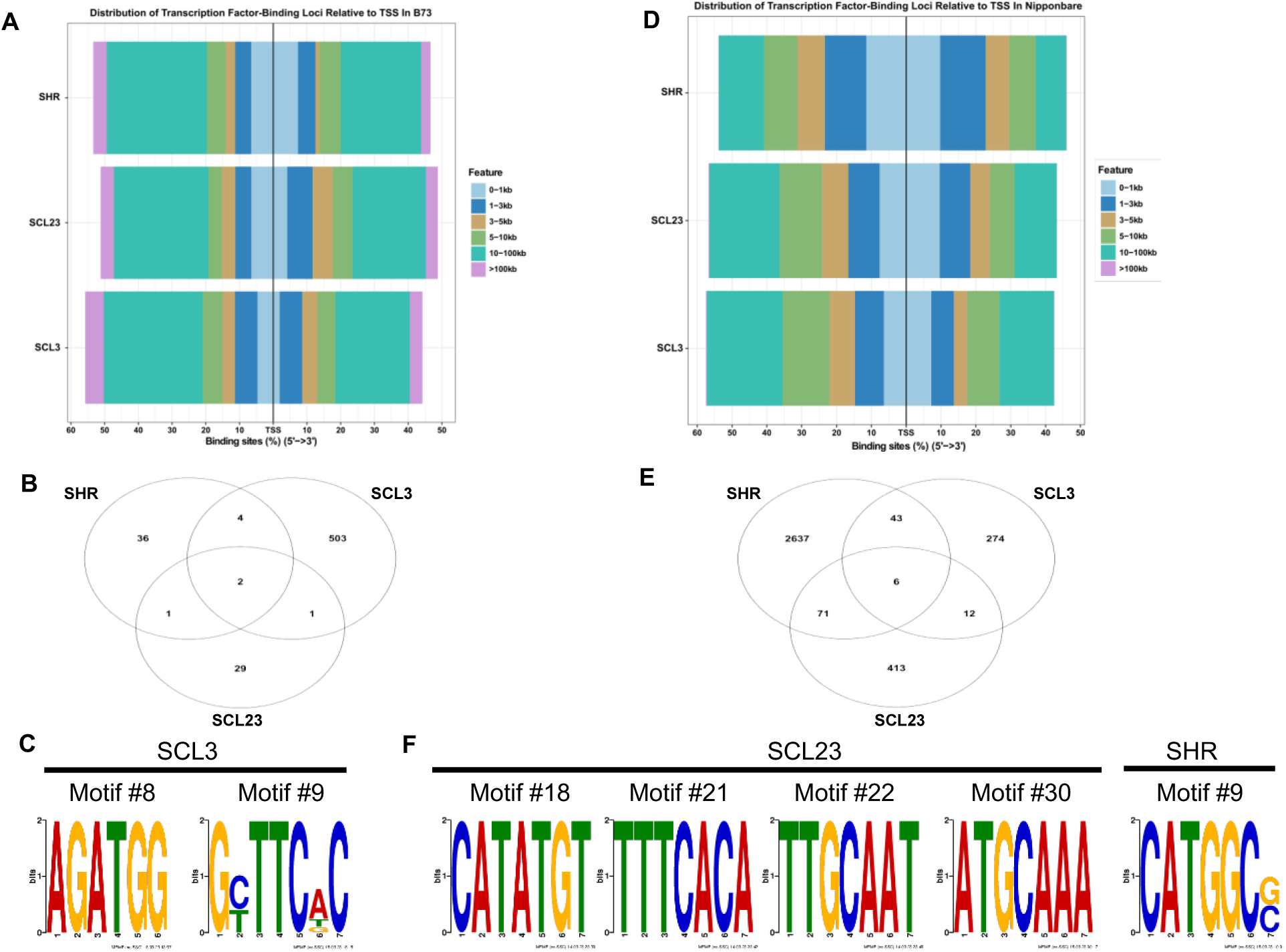
Sorghum GRAS family binding profiles in maize B73 and oryza Nipponbare. A) Distribution of all significant peaks for SbSHR, SbSCL23, and SbSCL3 relative to gene transcriptional start sites in the maize B73 genome. B) All maize B73 genes with GRAS peaks in their promoters. Called from the three DAP-seq pulldowns for Sorghum SbSHR, SbSCL23, and SbSCL3 using maize B73 DNA as the template. C) Uncharacterized DNA motifs that were detected in the SbSCL3 pulldowns in B73. D) Distribution of all significant peaks for SbSHR, SbSCL23, and SbSCL3 relative to gene transcriptional start sites in the Nipponbare genome. E) All rice Nipponbare genes with GRAS peaks in their promoters. Called from the three DAP-seq pulldowns for Sorghum SbSHR, SbSCL23, and SbSCL3 using Nipponbare DNA as the template. F) Uncharacterized DNA motifs that were detected in the SbSCL23 and SHR pulldowns in Nipponbare.

### Discovery of Novel DNA-binding Motifs in GRAS Family Transcription Factors

A DNA motif enrichment analysis was performed to determine what other TFs were active in the same regulatory space as SHR, SCL23, and SCL3. All GRAS peaks in the promoter region were analyzed using the MEME suite and several different classes of TF binding sites were identified as co-populating the peaks for either SHR, SCL23, or SCL3 (**Supplemental Figure 1 and Supplemental Data File 6**). The TF binding motifs observed to occur within the SHR promoter peaks included family members of AP2/EREBP, ABI3, TCP, NAC, and MYB (**Supplemental Figure 1a**). TF binding sequences found in SCL23 promoter peaks are AP2/EREBP, ABI3, GRF, DOF, Heat Shock Factor, bZIP, and MYB **(Supplemental Figure 1b**). For SCL3 promoter peaks, DNA-binding motifs were discovered for the TF families AP2/EREBP, ABI3, GARP G2-like, NAC, and MYB **(Supplemental Figure 1c**).

A subset of the enriched DNA motifs in the SHR and SCL23 peaks could be classified as novel GRAS motifs as they did not appear in existing TF-binding profile databases like JASPAR(Rauluseviciute et al., 2024) or Catalog of Inferred Sequence Binding Preferences (CIS-BP)(Weirauch et al., 2014). Additional bioinformatic analyses were conducted to determine if any of these putatively novel GRAS motifs were legitimate TF-binding sites due to the noisy nature of DAP-seq as a TF-profiling experiment. The position weight matrix (PWM) for each of the novel motifs discovered in SHR and SCL23 were projected across the promoter regions of all genes within sorghum (BTx623 v3), maize (B73 v5), and oryza (Nipponbare v1) genomes to determine their frequency of occurrence relative to the TSS (**Figure 3a**). The promoter region was binned into 40 bp intervals and each motif occurrence was counted. The motif sequence frequency in the promoter region, which we propose is unique to the three GRAS family TFs in this analysis, gradually reaches apogee at the −1000 bp position relative to the TSS. Then it decreases abruptly and then has a sharp, bimodal occurrence near the TSS (∼-400->+60 bp region), with some SCL23 profiles possibly displaying a 4th peak well into the coding sequence. These motifs had a unique frequency profile compared to other TF family DNA-binding motifs like WRKY, AP2/EREB, and bZIP proteins that can be more unimodal around the TSS but sometimes still display abrupt changes in frequency at different positions relative to the TSS **(Figure 3b).** There is variation of motif frequency conservation across sorghum, maize, and oryza for the SHR motifs; motif #29 shows fairly conserved frequencies across the three monocots, whereas SHR motif #26 shows strong frequency similarity between maize and sorghum, but the oryza profile seems to be ∼50% of that. The frequency of these novel, previously uncharacterized motifs from the maize and oryza DAP-seq analysis (previous section) also displayed similar frequency profiles to that of sorghum-derived pulldowns in oryza and sorghum genomes. All sorghum genes that have GRAS TF DAP-seq peaks as well as motif frequency occurrences from the PWM projections can be found in **Supplemental Table 4**. This suggests that 1) the gene modules that could be regulated by these TFs via their promoter regions are not completely conserved, 2) there is some incompleteness in the motif fingerprint projection, 3) GRAS TFs can recognize multiple promiscuous sequences, and/or 4) the motifs we identified only represent a portion of a larger recognition motif that is defined by *in vivo* binding activity. Ultimately, this analysis likely indicates that these previously uncharacterized motifs are more likely to be real DNA-binding sites due to their unique frequency and conserved occurrence around coding sequences in different monocots.

**Figure 3.**
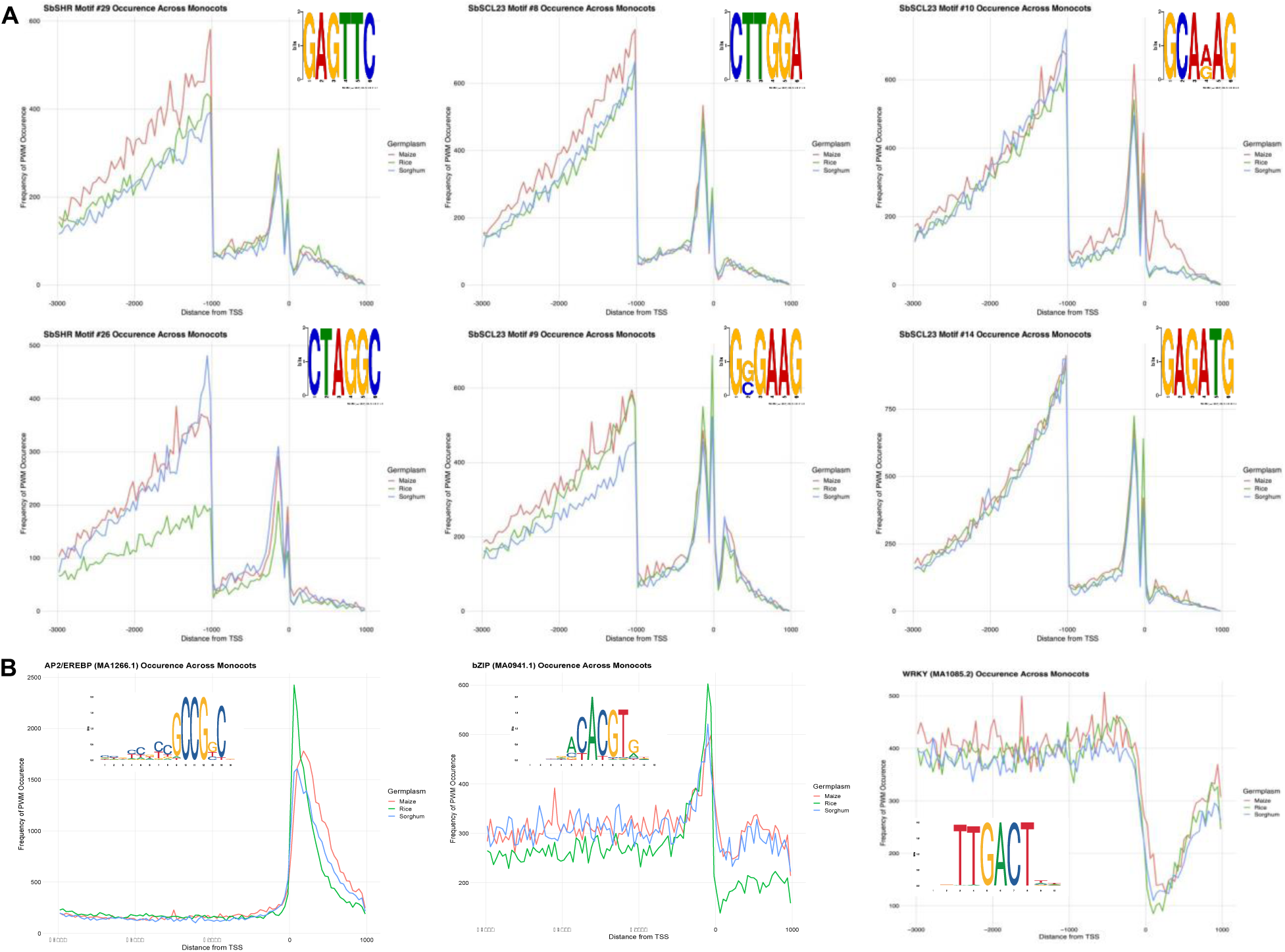
DAP-seq PWM projections in sorghum, maize, and oryza. A) The SHR motifs and SCL23 uncharacterized motifs identified in the BTx623 DAP-seq analysis. B) are some examples of common TF motifs projected across the genomes (taken from JASPAR).

### SCL23 Occupancy Around 3’UTR regions

While the focus of DNA-binding profiles tends to fixate around the promoters, TSS, and enhancer regions, TFs can bind in and around the 3’ untranslated regions (3’UTR) of gene models (**Supplemental data table 5**). The sorghum SHR, SCL23, and SCL3 TFs all displayed 3’UTR binding across hundreds of genes, often with no other nearby gene models. When evaluating DNA-recognition sites within 3’UTRs that host GRAS peaks, several environmentally responsive TF binding sites come out, specifically ARR-B/HHO, ABI3, and GRF motifs. There was no enrichment for the putative GRAS motifs identified within promoter peaks at 3’UTR regions. There are no consistent ontology enrichments between the three GRAS TFs for the genes with peaks in their 3’UTRs. However, when manually validating SCL23 peaks that 1) occur within 3’UTRs and 2) are not upstream of a neighboring gene TSS, numerous growth and developmental genes emerge. Some examples are SORBI_3001G261545, a DNA photolyase that is essential for phosphorus starvation response in roots (Nilsson et al., 2007); a strigolactone biosynthesis gene SORBI_3005G168200 that is strongly upregulated in response to limiting phosphorus conditions in roots(Gladman et al., 2022); SORBI_3002G075900, a dolichyl-diphosphooligosaccharide-protein glycosyltransferase protein whose orthologs plays a role in proper primary and lateral root formation in oryza(Qin et al., 2013), a cysteine desulfurase domain (SORBI_3002G174900) and a MATE family transporter (SORBI_3002G232600) that are strongly upregulated in roots in response to abiotic stress(McCormick et al., 2018; Gladman et al., 2022), and an exocyst complex component, SORBI_3003G158700, whose orthologs are involved in a variety of targeted cell secretions for proper cell progression and polarization(Pecenková et al., 2017; Synek et al., 2021).

### Shared Peaks Highlight Tissue-specific Gene Expression

Evaluating peaks that are shared between all three GRAS TFs revealed conserved promoter and enhancer elements within the sorghum BTx623 genome. Incorporating epigenetic information with the SHR, SCL23, and SCL3 binding locations also gives more power in identifying real versus spurious binding sites as well as yielding tissue-specific gene expression targets. To do this, all promoter DAP-seq peaks that have partial or near total overlap between SHR, SCL23, and SCL3 and were compared to Histone 3 trimethylation on the 4th lysine (H3K4me3) peaks that were derived from whole BTx623 roots grown in hydroponic conditions during a limiting phosphate experiment (data from Gladman, et al., 2022). H3K4me3 peaks generally indicate active gene expression and are often localized in a narrower fashion around the TSS, however H3K4me3 peaks can also be more broad and exist in the upstream promoter regions or distal enhancer space, and are likely indicative of tissue-specific expression in plants and other eukaryotes(Benayoun et al., 2014; Zhang et al., 2021).

Of the 256 overlapping peak regions that occur within promoters between all three GRAS TFs, 240 of them were associated with unique gene models (16 overlapping peaks occurred multiple times around the same gene element). Of the 240 unique shared peaks, 157 were manually confirmed to have near perfect overlap when visualized on a genome browser and 79 of those had almost complete overlap and exist on a region with notably higher H3K4me3 marks relative to the surrounding genome space (**Supplemental Data Table 6**). While gene expression doesn’t always correspond with upstream H3K4me3 peaks, 77 of the 257 overlapping peaks did show statistically different gene expression somewhere within the root system (data from Gladman, et al., 2022) during limiting phosphorus growth conditions. The majority of this significant differential gene expression occurred in the lateral root region, but there were also genes that showed differential expression in the root apex and elongation zone (**Figure 4a)**. Both up- and down-regulated genes were downstream of these strong GRAS TF peaks, so the presence alone of these TFs and H3K4me3 marks do not fully explain the regulatory nature of the native DNA binding capability of GRAS family proteins during nutrient stress response.

**Figure 4.**
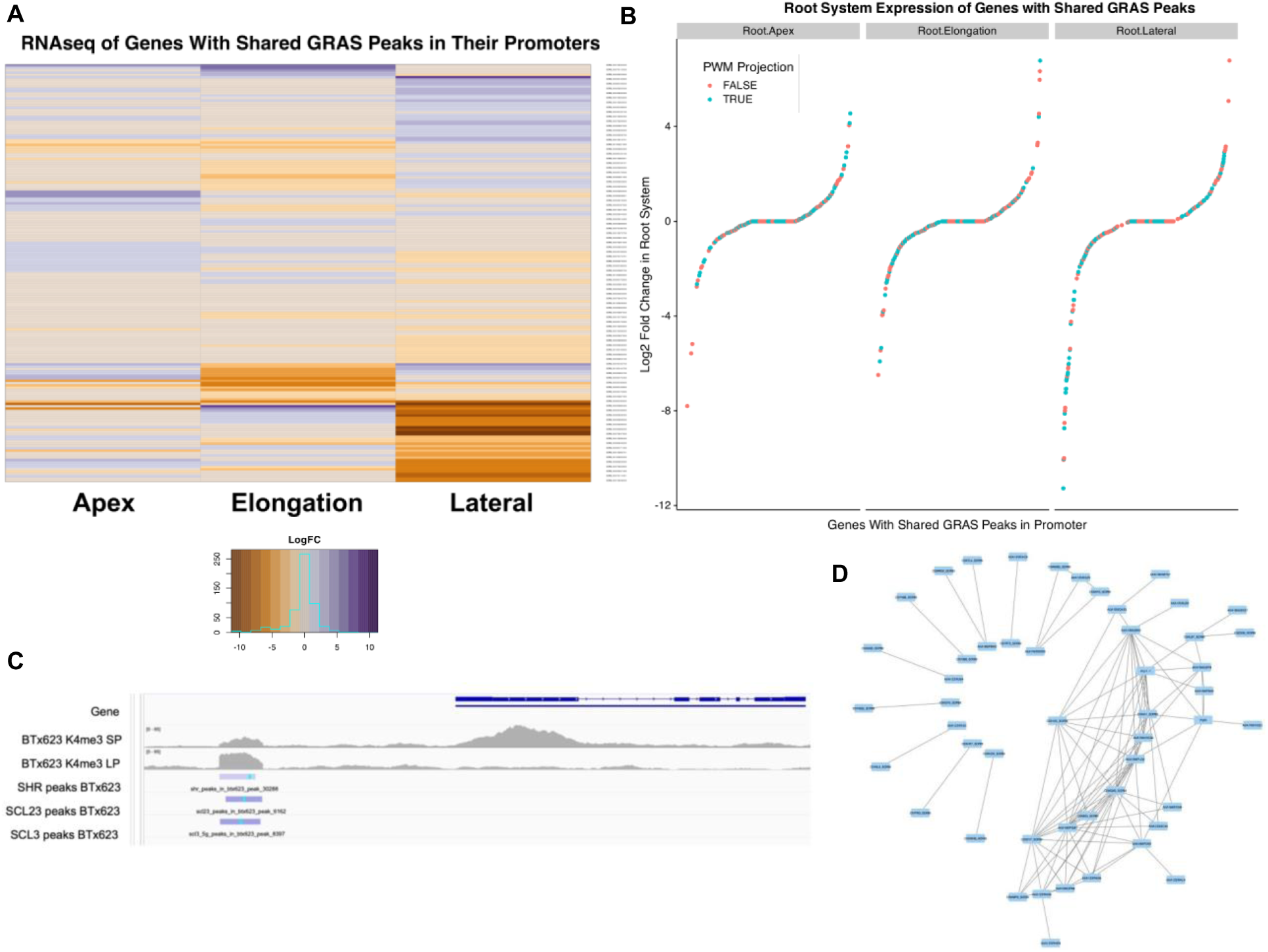
Expression and network analysis of the 240 genes whose promoters have shared binding peaks from SHR, SCL23 and SLC3. A) Log2-fold expression heatmap of genes that have peaks in their promoter region from all three SHR, SCL23, and SCL3. Tissue expression data is from the root apex, elongation zone, and lateral root regions (data from Gladman, *et al*. 2022). B) Log2-fold expression of (A) with colorized indication if those genes also were identified from the PWM projection analysis. C). Integrated Genome Viewer display of a LysM gene (SORBI_3002G222500) that has strong overlap between SHR, SCL23, and SCL3 and the localization of all three peaks corresponds with H3Kme3 pileup taken from whole root samples grown in hydroponics under normal and limiting phosphorus conditions (data from Gladman, *et al*. 2022). D) Cytoscape network display of available protein-protein interaction and co-expression data from the STRING-DB resource. The larger network on the lower right is enriched basic molecular ontologies (purine biosynthesis and translation).

To assess the accuracy of the PWM projections on these shared peaks, 186 of the 256 genes (72.7%) were present in at least one of the PWM projected gene groups from the above analysis (**Figure 4b**). Many of the genes that had near perfect overlap with all GRAS peaks; H3K4me3 marks; significant differential gene expression in the lateral, apex, or elongation root region; and existed in the PWM projections, are involved in cell growth, development, signaling, and environmental response. Some examples are the chromatin remodeler SORBI_3006G038400; the fasciclin-like arabinogalactan protein SORBI_3004G137200; the ABA/WDS protein SORBI_3008G049200 that is induced by water/ABA stress in oryza(Li et al., 2017); and the LysM domain-coding protein SORBI_3002G222500 (**Figure 4c**), which is involved in arbuscular mycorrhizal symbiosis(Yu et al., 2023). SORBI_3005G053501, a defense response gene, shows a greater H3K4me3 signal and is more strongly expressed in the root apex and elongation zone during sufficient phosphorus conditions. SORBI_3002G076100, a G-box TF that has been identified as playing roles in both photomorphogenesis with HY5(Singh et al., 2012) and also root hair development in Arabidopsis(Richter et al., 2011) has shared promoter binding between all three GRAS TFs and was shown to be upregulated in sorghum lateral root regions in response to limiting phosphorus(Gladman et al., 2022). When comparing the gene promoters that contained shared peaks with the PWM projection data on the putative SHR and SCL23-specific motifs, 50 out of 256 promoters (19.5%) had hits for at least one of the two SHR motifs and 156 out of the 256 promoters (49.2%) had hits for at least one of the four SCL23 motifs. Incorporating data from the STRING database, a network could be constructed based on existing protein-protein and co-expression information as well, which resulted in a smaller network that was statistically enriched for purine biosynthesis (GO:0009205 and GO:0006164) and translation (GO:0006412) (**Figure 4d and Supplemental Data Table 7**).

Additionally, this multi-omics integration identified multiple short gene models that have no current functional domain annotation (e.g. SORBI_3007G226900, SORBI_3010G238300, SORBI_3005G145800, SORBI_3010G201332, SORBI_3002G149600, SORBI_3006G024450), yet have significant expression in root tissue or abundant H3K4me3 marks with the trio of GRAS peaks in the promoter, or both. This provides evidence that these are real gene models and likely active in the root transcriptome (**Supplemental Figure 2**). Importantly, when comparing DAP-seq peaks with the conserved cis-regulatory element information from the Conservatory Project(Hendelman et al., 2021), the GRAS-specific regulatory regions did not usually coincide with an evolutionarily constrained cis-regulatory elements, suggesting that while GRAS TFs are quite old in plants, their regulatory sites can undergo re-wiring in a species-specific manner.

## Discussion

Prior experimentation on GRAS DNA-binding is scant, especially considering the importance of the family to multiple functions across plant systems. Our DAP-seq profiling demonstrated good agreement with other work in *Arabidopsis*, and targeted genes yielded functional ontologies that could correspond with prior genetic and molecular characterizations of *SHR* and *SCL23* in other species. Specifically, all three GRAS TFs displayed gene target ontology associations for previously characterized functions like flavonoid biosynthesis(Pillet et al., 2015; Huang et al., 2021) and as well as basal molecular functions, especially with essential cofactor metabolism, cellular respiration, and cell wall/membrane-associated processes. This broad involvement of genetic function for these TFs could reflect the long co-evolution of GRAS genes since they first emerged in pre-vascularized plants.

TF profiling in combination with identifying conserved non-coding sequences has become a powerful method to identify targets for gene editing approaches(Rodriguez-Leal et al., 2019; Hendelman et al., 2021; Liu et al., 2021a; Aguirre et al., 2023) as well as improve gene regulatory networks inference through machine learning approaches(Shojaee and Huang, 2023). Despite DAP-seq being a ‘noisy’ assay to profile TF binding sites within a genome, it provides a notable benefit over similar assays like ChIP-seq and CUT&RUN: cell-type naive binding. While DAP-seq profiling precludes binding behavior of TFs that require or are modified by cell-type specific cofactors and DNA and histone methylation, native DNA binding behavior of TFs can be assessed and winnowed for high-confidence peaks through the inclusion of tissue-specific transcriptomic and epigenetic data. This creates the capacity to generate GRNs across multiple tissues by performing DAP-seq once and then layering that data with tissue-specific gene expression information and epigenetic marks. We have used this paradigm in our novel characterization of the important plant-specific GRAS family of TFs and combine additional expression and epigenetic data to improve the confidence in calling biologically significant binding sites for SHR, SCL23, and SCL3 in *Sorghum bicolor*.

Confirming the presence of TF binding motifs is more straightforward when working with TFs that have previously characterized binding sequences, such as WRKYs, NACs, bHLH, and others. It is hard to confidently assert that a previously uncharacterized motif sequence is a true TF binding recognition site with DAP-seq data. However, we found that filtering for SHR and SCL23 peaks, for example, that occur in the promoter regions of genes, then evaluating the frequency of those motifs across the promoters of the entire genomes for sorghum, oryza, and maize created additional supportive evidence that 1) those DNA-binding motifs are likely real and 2) they are likely recognized by the GRAS TFs due to the unique pattern of their occurrence relative to other well characterized TFs. This type of bioinformatic approach allows for additional means of identifying putative regulatory targets for TFs that can bind to projected motifs as well as provide insight into the evolutionary conservation of them across species.

Our first-ever DAP-seq profiling of the sorghum SHR, SCL23, and SCL3 proteins using B73 and Nipponbare input DNA yielded significantly fewer peaks in both maize and oryza compared to the original sorghum gDNA template. This disparity in peak enrichment using the SbGRAS proteins could be due to the divergence in protein sequence identity between the closest orthologs of SbSHR, SbSCL23, and SbSCL3 in oryza and maize. Another explanation for this peak enrichment disparity is there is significant rewiring in the promoter and enhancer space of recognition sites for these three GRAS family motifs. However, both of these explanations rely upon the non-conservation of the regulatory motifs across these three species, which confounds our motif occurrence projections as described above and in **Figure 3**. Ultimately, this could reflect that the putative GRAS-specific motifs we identified for SbSHR and SbSCL23 are only representative of a part of a larger cis-regulatory element sequence that cannot be resolved through the MEME suite program using DAP-seq peak inputs. There is evidence that single DAP-seq might only capture a small window of a larger biologically active binding site for TFs that require complexing with other TFs or cofactors(Li et al., 2023). This hypothesis could absolutely apply to the GRAS TFs, and SHR and SCL23 in particular, since it is well known that they interact with other TFs and protein cofactors as well as each other to heterodimerize and modulate binding functions in the nucleus.

Ultimately, the power of this research lies in the GRNs that are generated. They reveal potential GRAS gene targets that can be leveraged by and for breeding programs and functional research through identification of disruptive alleles and CRISPR editing approaches. Furthermore, this analysis affirms that GRAS TFs do have innate DNA-binding activity without interacting with other protein cofactors or TFs like IDDs. Identifying these regulatory upstream promoter sequences are useful for EMS mutagenesis or CRISPR genome editing approaches to fine tune gene expression for a spectrum of agronomically valuable phenotypes. Additionally, these types of profiling experiments can also reveal tissue-specific regulatory DNA regions that are being acted upon by promiscuous TFs families, which could permit more precise genome editing that could impact intended organs and not have systemic effects across the plant. For example, SHR has been shown to be involved in arbuscular mycorrhizal symbiosis, and indeed through our GRNs and PWM projections, we show that all three GRAS TFs bind to a very narrow promoter region upstream of a gene involved in arbuscular mycorrhizal symbiosis, LysM (SORBI_3002G222500). This promoter region also displays strong epigenetic signals for open chromatin; yielding a strong candidate for EMS mutagenesis or CRISPR genome editing for those interested in nutrient use efficiency. Ultimately, this first-ever GRAS profiling in sorghum combined with additional -omics data has generated a list of useful targets for additional agronomic characterization that span everything from nutrient use efficiency to growth and developmental modification.

## Methods

### DAP-seq Pulldown and Sequencing

DAP-seq was performed in a modified fashion from the original methods(O’Malley et al., 2016). *Scarecrow Like 3 (SCL3; SORBI_3005G029600), Scarecrow Like 23 (SCL23; SORBI_3002G342800),* and *Short Hair Root (SHR; SORBI_3001G327900)* full length coding sequences (CDSs) were synthesized from TwistBiosciences (pTwist-ENTR Kozak vector) and cloned into the pDEST15 Gateway vector (N-terminal GST-tag). The resulting plasmids were transformed into BL21 competent cells. Expression of GST-tagged TF proteins was induced between OD_600_=0.500-0.600 by the addition of 1mM isopropyl-beta-D-thiogalactoside (IPTG) (Goldbio: I2481C25) and 0.05M or 0.1M D-mannitol to bacteria in Lysogeny Broth. A concentration of 0.05M D-mannitol was added to SCL3 expressing bacteria, and 0.1M D-mannitol was added to both SCL23 and SHR expressing bacteria. The bacteria were then shaken at 16°C at 220 rpm for 16 hours. Cultures were spun down and pellets resuspended in 0.5M D-mannitol PBS. Cell membranes and plasmid DNA were disrupted by sonicating at 4°C in 10 cycles of 30 sec on, 30 sec off. Soluble fractions of lysate were added to triplicates of MagneGST beads (Promega) suspended in equal volumes of 0.5M D-mannitol PBS and incubated at 4°C for 1.5hrs undergoing end-over-end mixing to bind GST-tagged transcription factor proteins. The protein-bound beads were then incubated in sheared adaptor-ligated high-purity, sheared genomic DNA of sorghum BTx623, oryza Nipponbare, or maize B73, using between 100-250 ng of genomic DNA per sample.

Genomic DNA from these plants were sheared by Covaris S220 sonicator. Adaptors were ligated to genomic DNA fragments according to NEBNext Ultra II DNA Library Prep Kit and size-selected using AMPure XP beads for fragments larger than 300bp. Transcription factor-bound DNA fragments were then washed three times with the mannitol PBS solution and eluted off the beads by incubating at 98℃ for 5 minutes. qPCR was performed to determine the number of cycles needed during PCR to amplify eluted fragments and add barcodes; this PCR was performed with both elutes and fragmented input DNA as control, using 10-18 cycles depending on the library. PCR products were cleaned up using AMPure XP magnetic beads. Concentration was determined by Qubit HS dsDNA kit. Quality check was performed and average size of the libraries determined by Bioanalyzer on High Sensitivity DNA Chips. qPCR was performed to determine concentration of the barcoded libraries. Samples were pooled and sent for sequencing. *S*equencing was performed at paired end 150 high output using the Illumina NextSeq2000 platform for the sorghum BTx623 samples and on the Illumina Novaseq 6000 platform for the maize B73 and oryza Nipponbare samples. All 36 libraries (3 replicates of SHR, SCL23, and SCL3 pulldowns + Input control) from each sorghum, oryza and maize were multiplexed into 3 total pools (12 libraries per pool), yielding between 22.1-66.3 million reads per sample for the *Sorghum bicolor* DNA query (BTx623), 9.7-27.2 million reads per sample for the *Zea mays* query (B73), and 8.9-33.8 million reads per sample for the *Oryza sativa (*Nipponbare) query.

### DAP-seq Bioinformatic Analysis

The FASTQ files were aligned and merged as follows: Trimmomatic(Bolger et al., 2014) was used for FASTQ trimming, followed by BWA mem alignment to the BTx623 v3 genome and MACS3 callpeak (v3.0.0b1) peak calling (using the input controls for background subtraction), and finally the annotatePeaks program from the ChipSeeker package(Wang et al., 2022) was used to associate peaks with gene models from the reference genome files. For ChipSeeker, default values for intergenic space was defined was used >10,000 bp upstream or >300 bp downstream of gene elements. Promoter regions were defined as <2,000 bp from the TSS.

The *Sorghum bicolor* BTx623 v3 reference genome files housed by Gramene(Release 66)(Tello-Ruiz et al., 2022) were used for annotating peaks. Sorghum GFF and GTF files were both used for Chipseeker features functionality; SAMtools was used for various file formatting and manipulation steps, including sorting and merging of the 150-bp paired-end read files. The version 1 *Oryza sativa* Nipponbare reference genome files housed by Gramene (Release 66) (Tello-Ruiz et al., 2022) were used for oryza mapping and peak calling. The version v5 *Zea mays* B73 reference genome files housed by Gramene (Release 66)(Tello-Ruiz et al., 2022) were used for maize mapping and peak calling. Motifs were compared between the results of each genomic DNA background. Motif enrichment analysis was performed using the MEME suite(Bailey et al., 2015). Identification of DNA-recognition motifs were done by comparing DAP-seq PWMs to the JASPAR nonredundant plant database(Rauluseviciute et al., 2024). Gene ontology analysis was done by submitting genes to the Gene Ontology Consortium online tool (https://geneontology.org/)(Ashburner et al., 2000) and only the Fisher’s Exact statistical test was used to calculate for significant enrichment without any correction.

### Cis-regulatory Motif Frequency Projections

To determine cis-regulatory elements for GRAS TFs within the promoters of sorghum, maize and oryza, we used a previously described computational prediction pipeline(Eveland et al., 2014; Knauer et al., 2019) (Eveland et al, 2014; Knauer et al 2019) that uses the Search Tool for Occurrence of Regulatory Motifs (STORM) from the Comprehensive Regulatory Element Analysis and Detection (CREAD) suite of tools (Smith et al., 2006; Schones et al., 2007). We identified GRAS PWMs from the MEME analysis (see above) and used these PWMs to identify motifs within the promoter region spanning 3kb upstream and 1kb downstream from the transcription start sites of all of the protein coding genes of sorghum BTx623, maize b73 and oryza Nipponbare. We considered only those motifs that were over-represented (*p*-value <0.001) in promoter sequences of protein coding genes as compared with a background set of the same number of random genomic sequences and these predictions were further filtered based on a PWM-specific median score threshold (i.e., quality score greater than or equal to the median score passed the filter) and a motif occurrence frequency of two or more per promoter.

## Supporting information

Supplemental_data_file_1_shr_peaks_sorghum_BTx623

Supplemental_data_file_2_scl23_peaks_sorghum_BTx623

Supplemental_data_file_3_scl3_5g_peaks_sorghum_BTx623

Supplemental_data_file_4_b73_3kb_to_1kb_downstream_tss_meme_output

Supplemental_data_file_5_nipponbare_3kb_to_1kb_downstream_tss_meme_output

Supplemental_data_file_6_Btx623_3kb_to_1kb_downstream_tss_meme_output

## Data Availability Statement

The DAPseq reads are available at the NCBI SRA repository, BioProject ID = PRJNA1162020.

## Author Contributions

NG and DW experimental design. NG performed the peak calling, annotation, and motif analysis. SK performed the PWM frequency projection analysis. AF and MR performed the DNA template preparation and DAP-seq pulldowns. All authors contributed to the manuscript. No AI/LLM was used in the writing of this manuscript.

## Acknowledgements

The authors would like to thank members of the plant research community at Cold Spring Harbor Laboratory for their feedback and support. This work was performed with assistance from the US National Institutes of Health Grant S10OD028632-01. This project was funded by the USDA-ARS award number 8062-21000-044-000D and 8062-21000-051-000D.

---

**Supplemental figure 1. DNA binding motifs in GRAS transcription factor peaks.** These are the DNA recognition motifs that were enriched under the DAP-seq peaks within gene promoters for A) SHR, B) SCL23, and C) SLC3

**Supplemental Figure 2.**
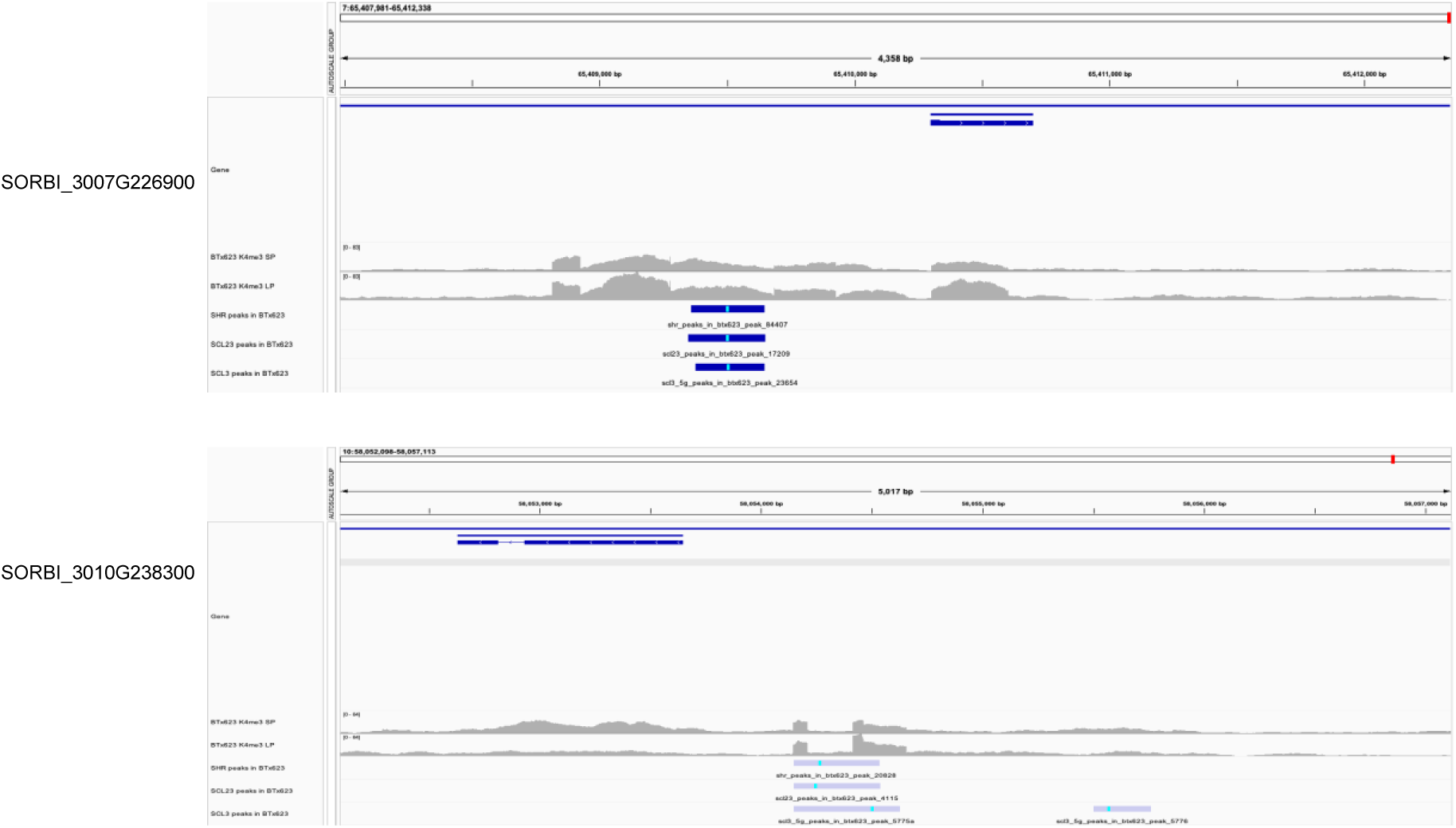
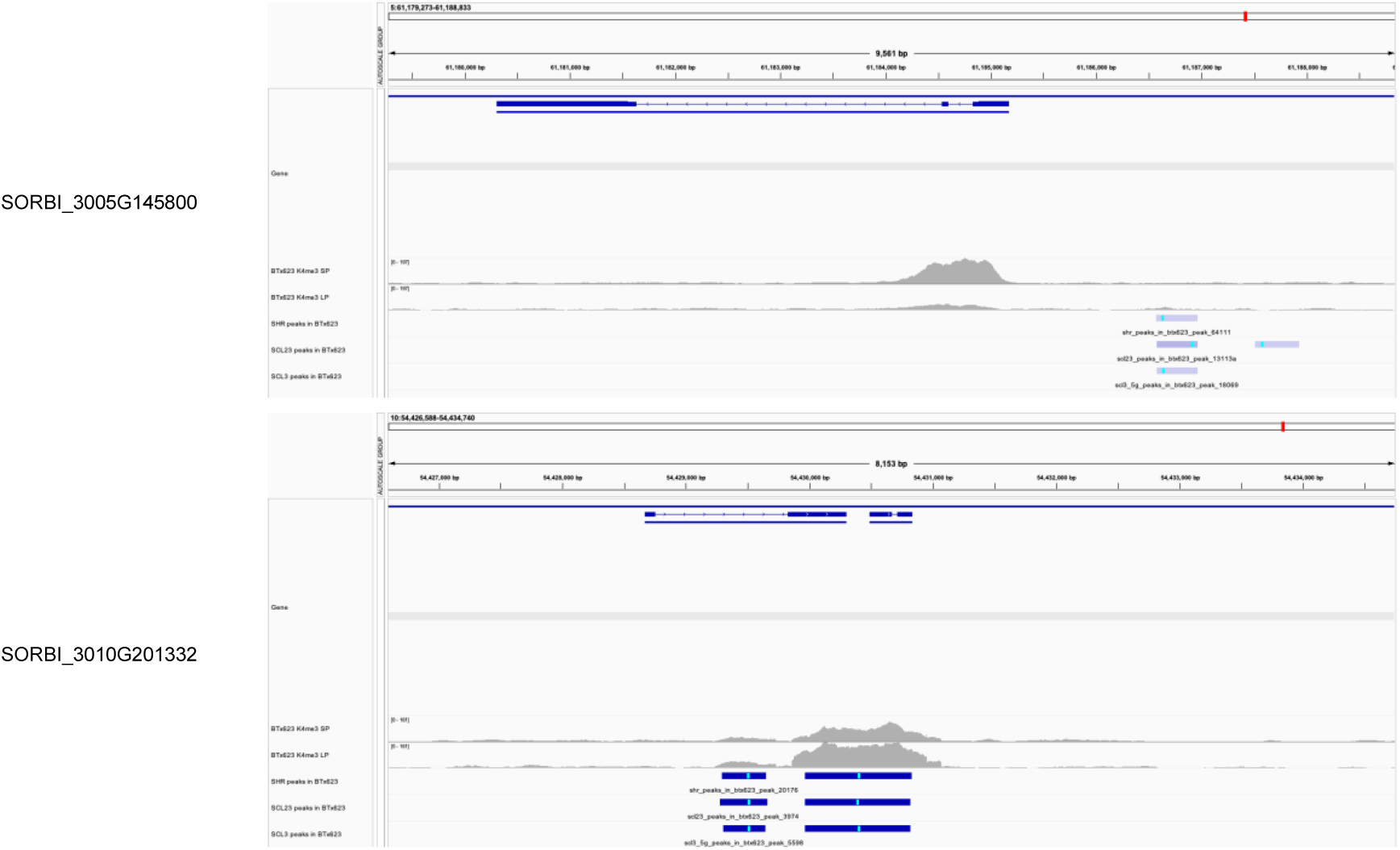
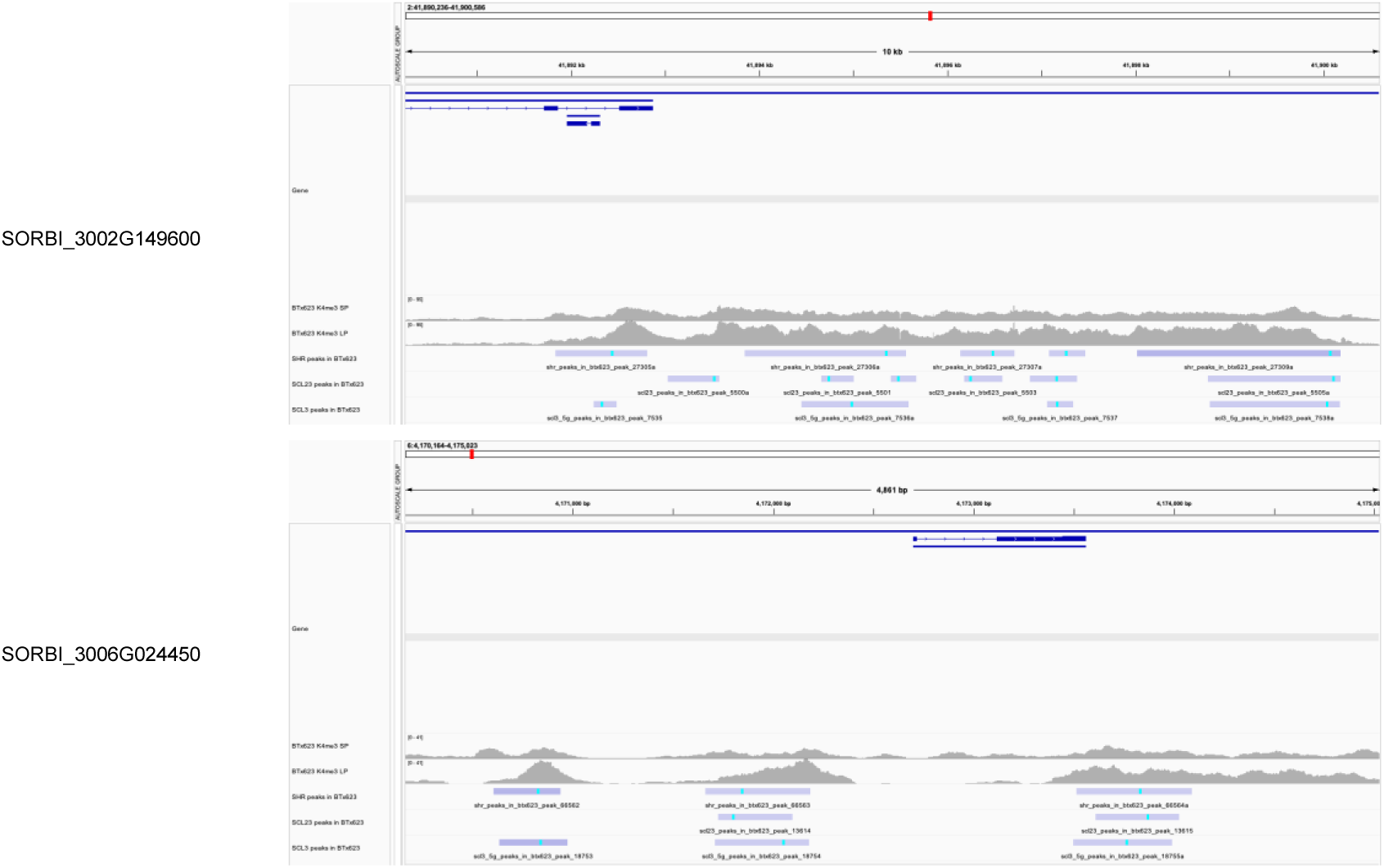
Combining GRAS DAP-seq peaks and root H3K4me3 marks. Integrated Genome Viewer images of examples of short genes (protein polypeptide product < 50 amino acids) that have overlapping GRAS family DAP-seq peaks and H3Kme3 pileups in the promoter region. (Histone methylation data taken from Gladman *et al*., 2022).

