## Supplementary figures and images for "GRAS Family Transcription Factor Binding Behaviors in Sorghum bicolor, Oyrza, and Maize"

### logo2.png

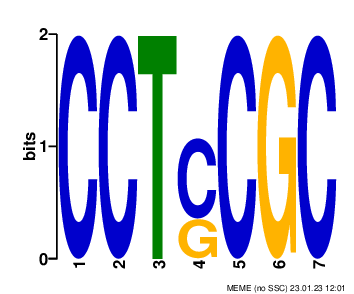

### logo4.png

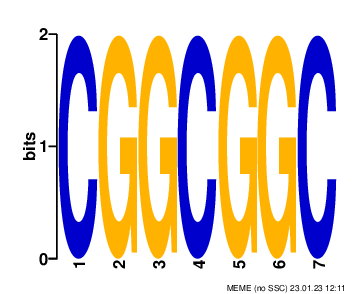

### logo5.png

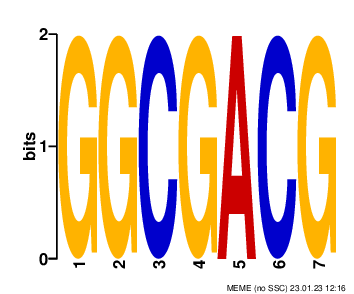

### logo6.png

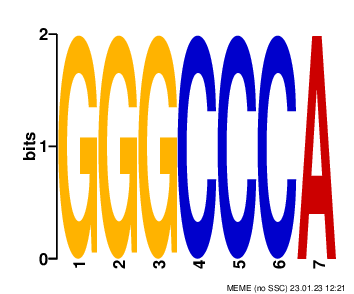

### logo7.png

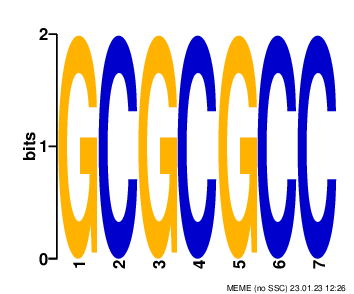

### logo20.png

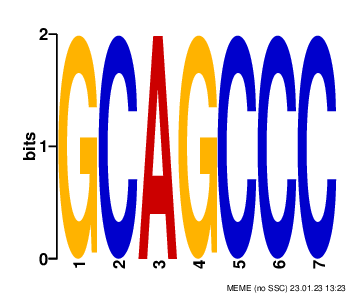

### logo21.png

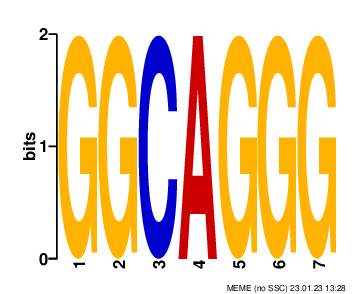

### logo22.png

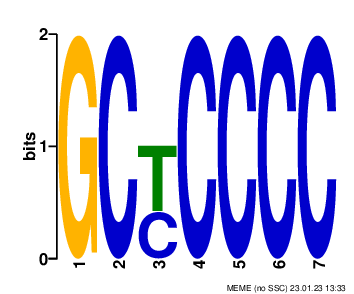

### logo23.png

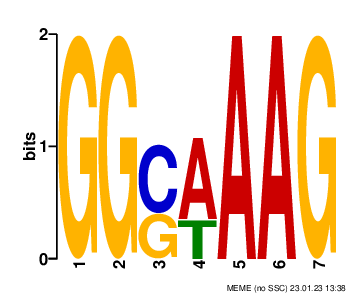

### logo25.png

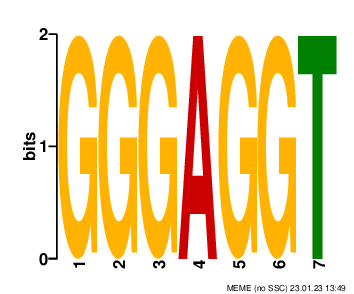

### logo_rc4.png

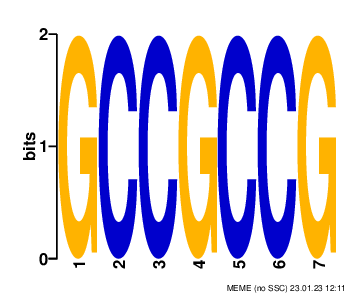

### logo_rc5.png

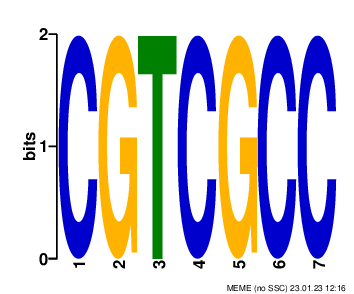

### logo_rc6.png

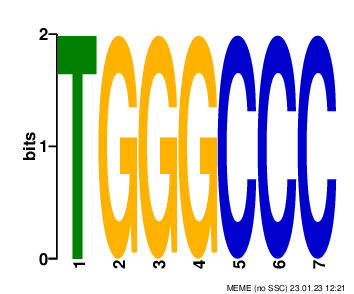

### logo_rc7.png

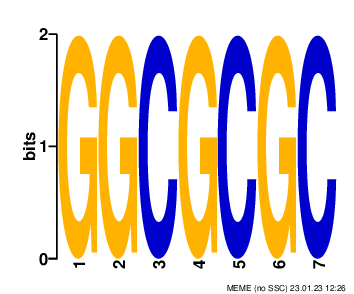

### logo_rc18.png

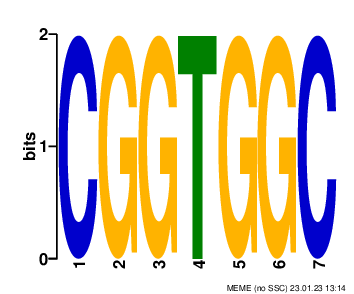

### logo_rc19.png

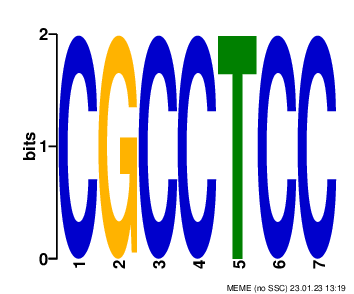

### logo_rc24.png

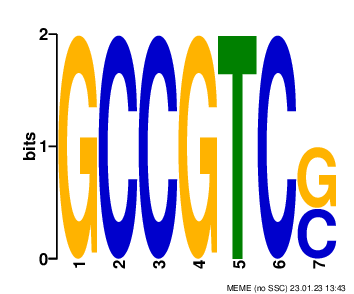

### logo_rc25.png

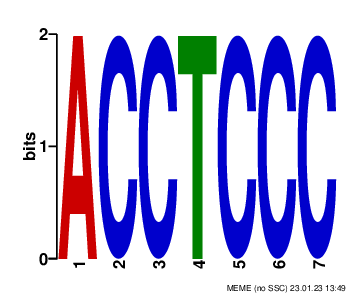

### logo_rc26.png

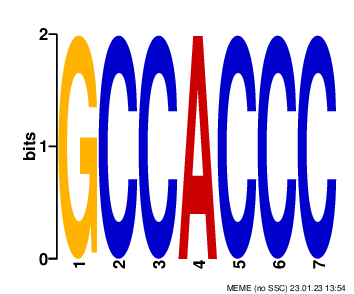

### logo_rc27.png

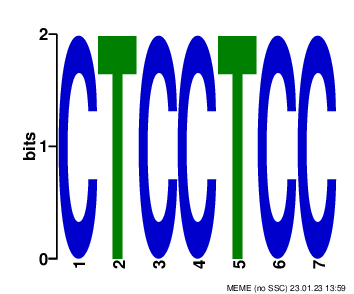

### logo_rc30.png

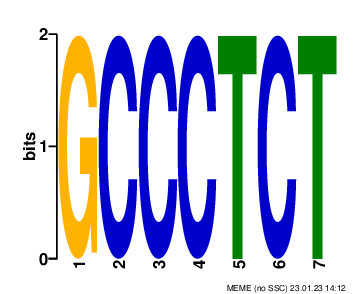
