## Supplementary figures and images for "GRAS Family Transcription Factor Binding Behaviors in Sorghum bicolor, Oyrza, and Maize"

### logo1.png

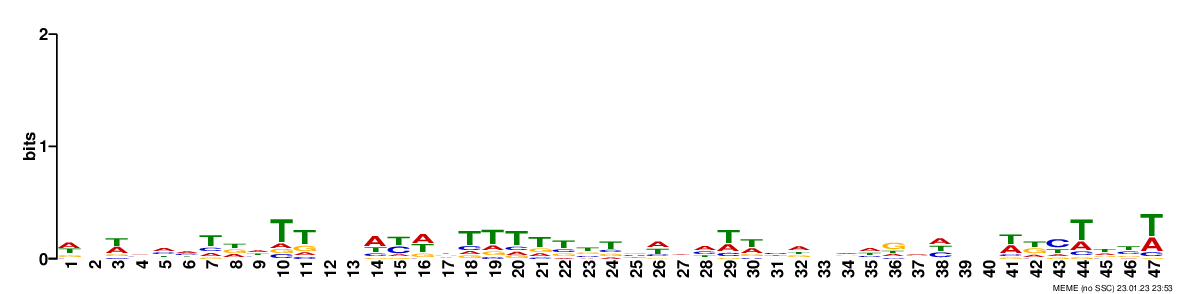

### logo1.png

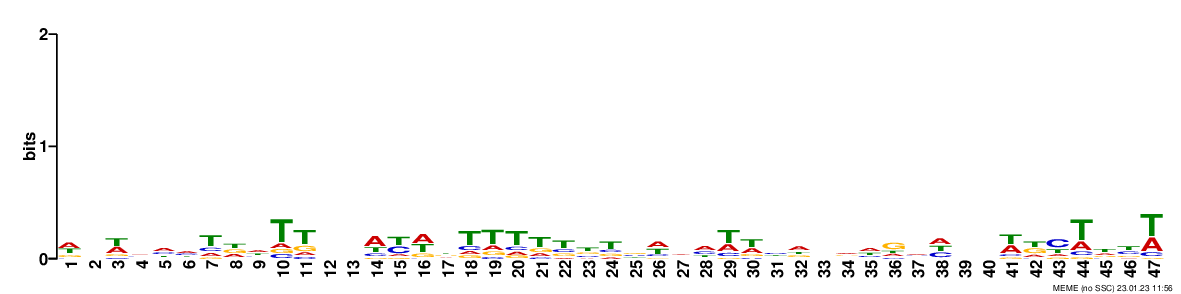

### logo2.png

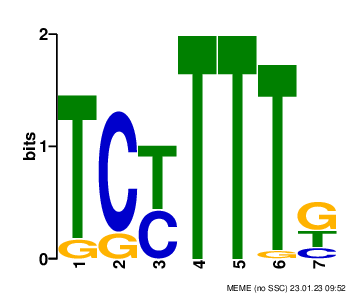

### logo2.png

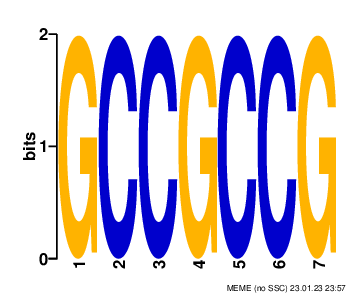

### logo3.png

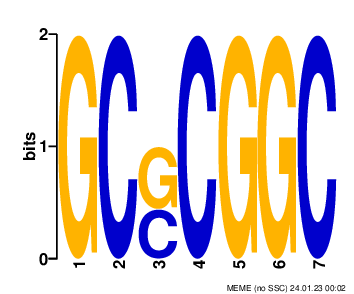

### logo3.png

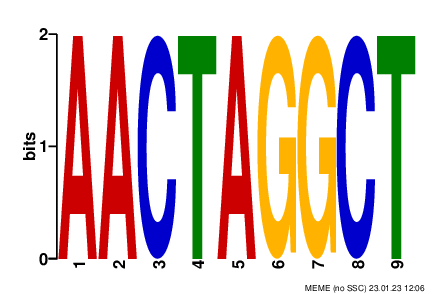

### logo4.png

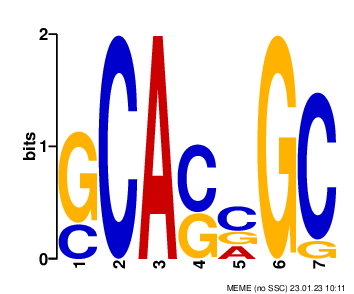

### logo4.png

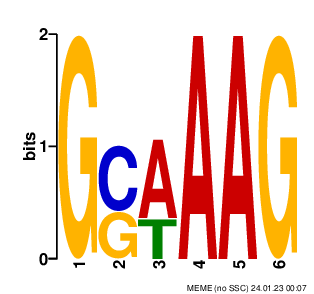

### logo5.png

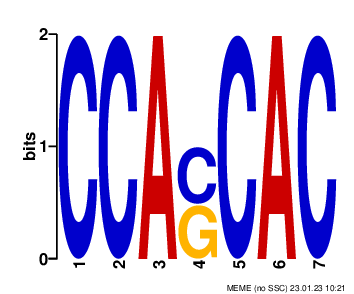

### logo5.png

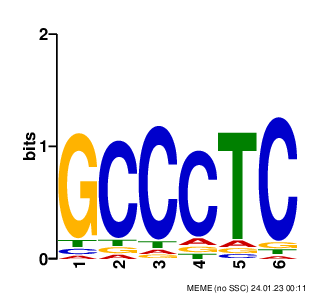

### logo6.png

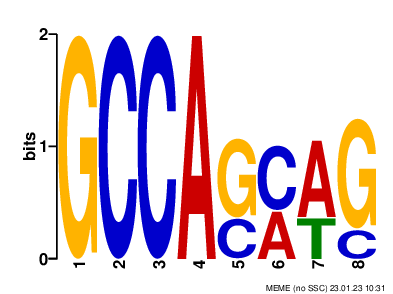

### logo6.png

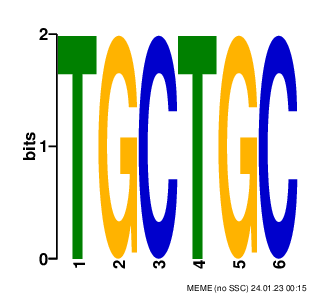

### logo7.png

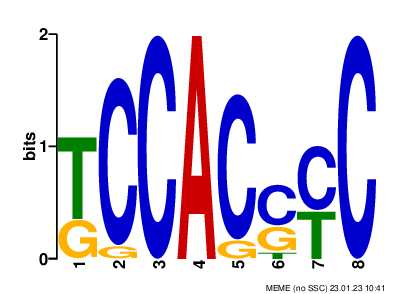

### logo7.png

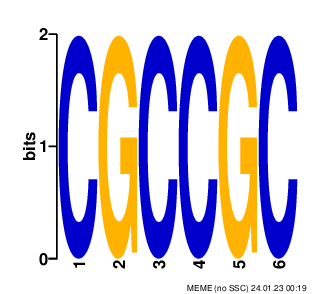

### logo8.png

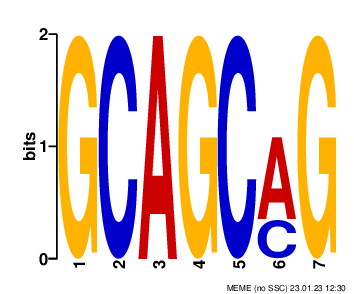

### logo9.png

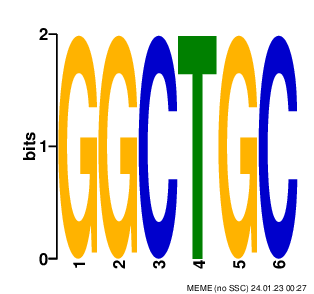

### logo9.png

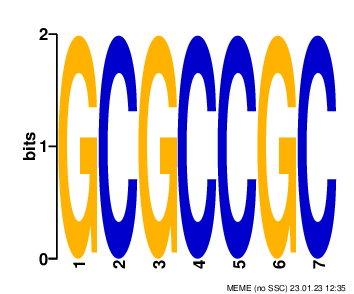

### logo10.png

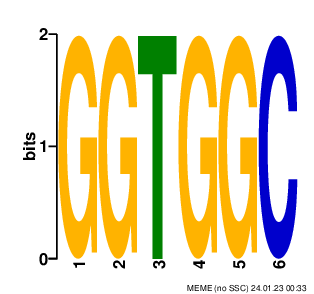

### logo10.png

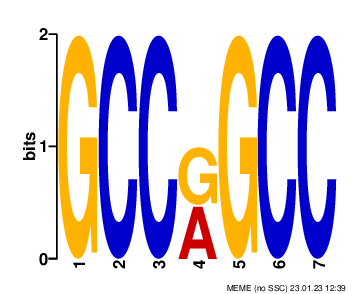

### logo11.png

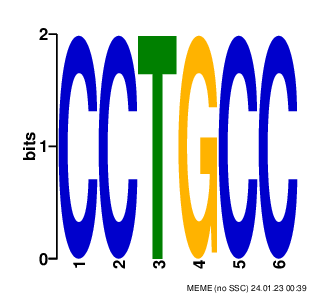

### logo11.png

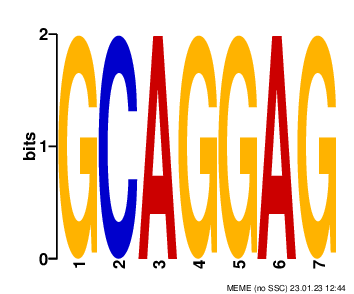

### logo12.png

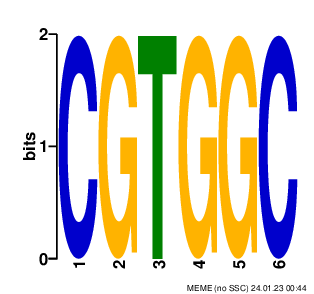

### logo12.png

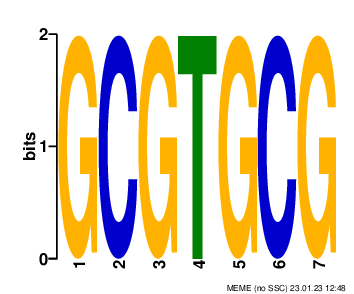

### logo13.png

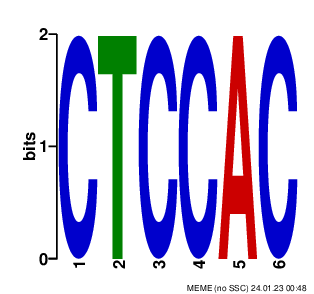

### logo13.png

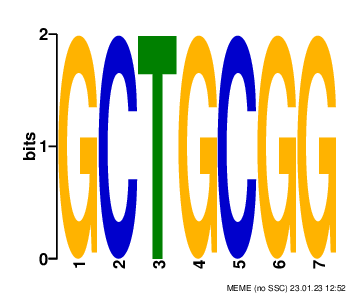

### logo14.png

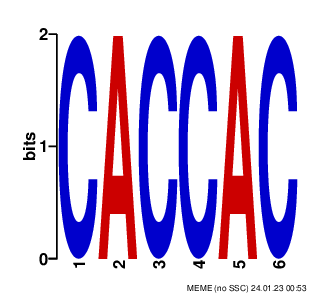

### logo14.png

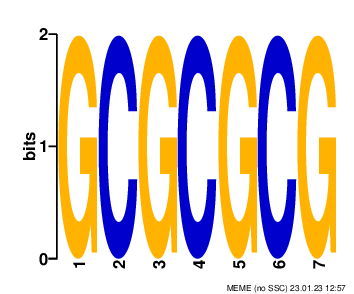

### logo15.png

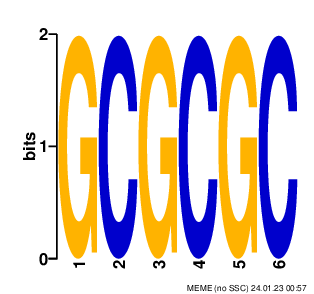

### logo15.png

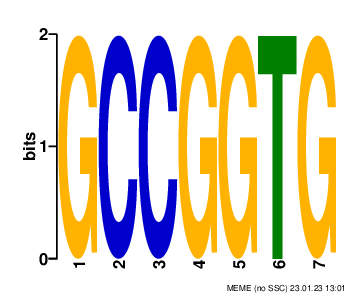

### logo16.png

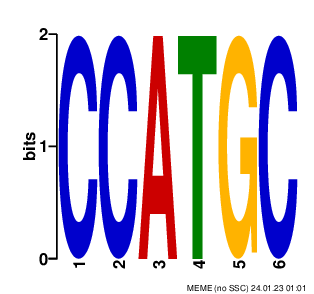

### logo16.png

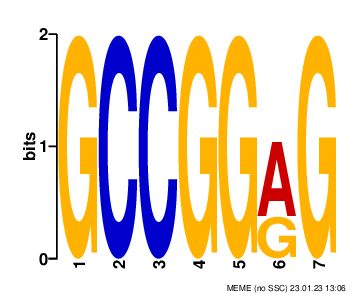

### logo17.png

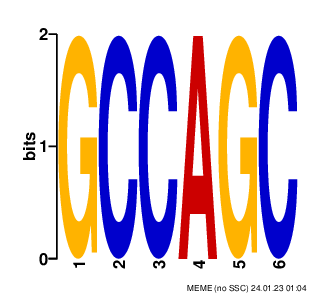

### logo17.png

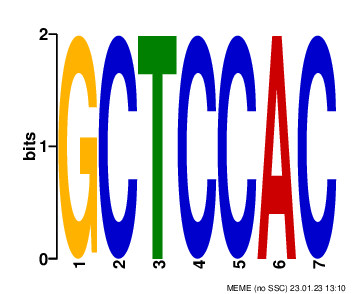

### logo18.png

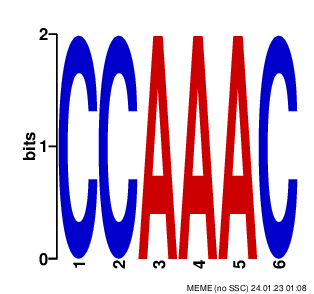

### logo19.png

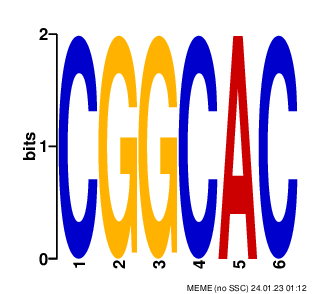

### logo20.png

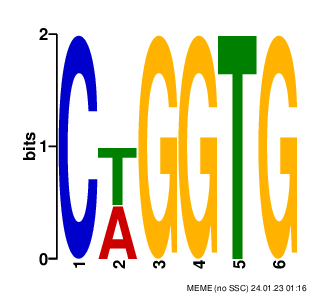

### logo21.png

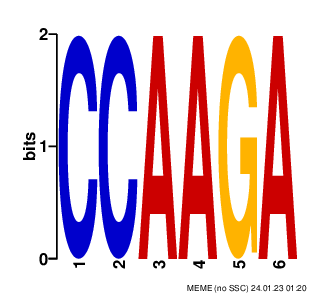

### logo22.png

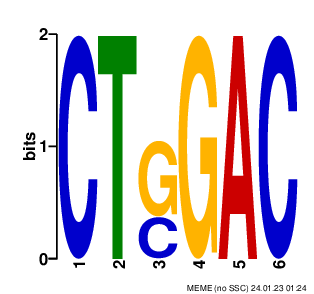

### logo23.png

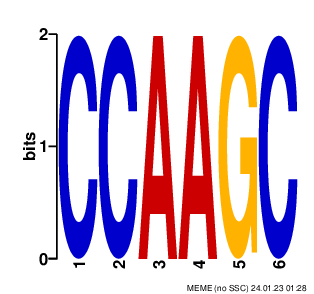

### logo24.png

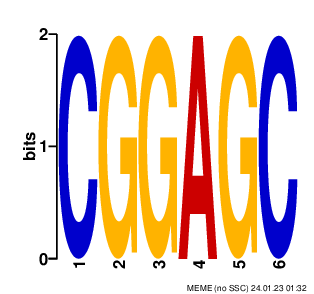

### logo25.png

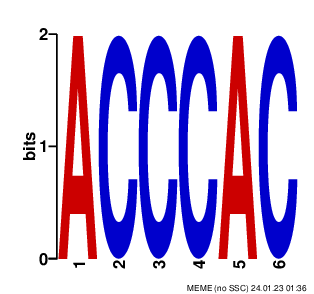

### logo26.png

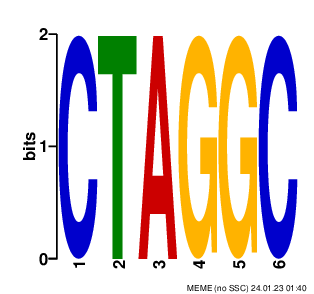

### logo27.png

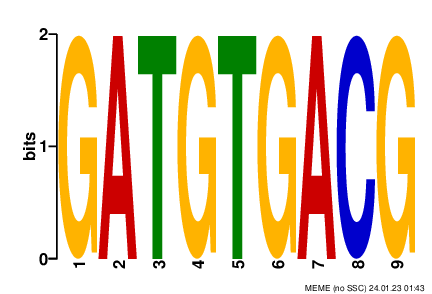

### logo28.png

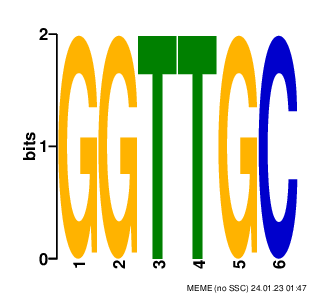

### logo28.png

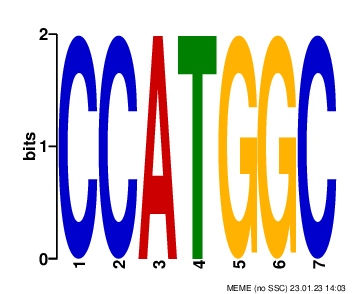

### logo29.png

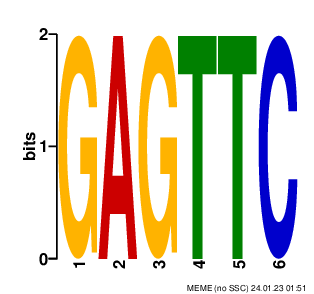

### logo29.png

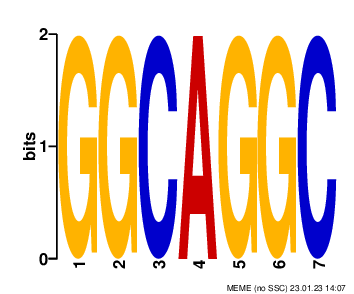

### logo30.png

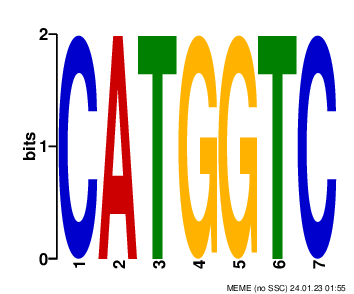

### logo_rc1.png

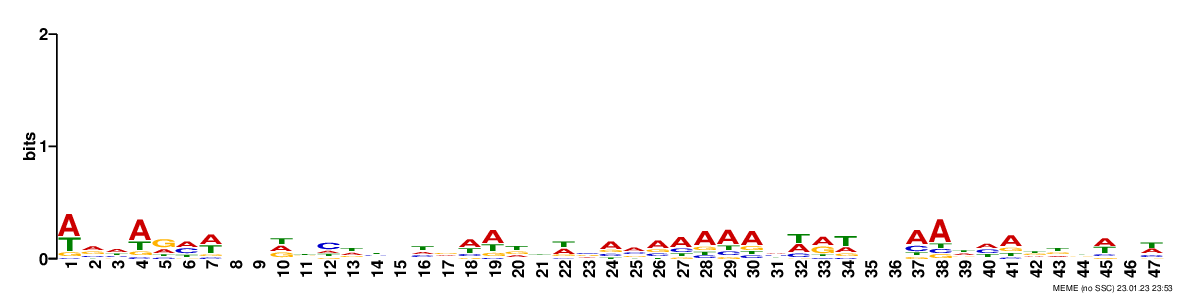

### logo_rc1.png

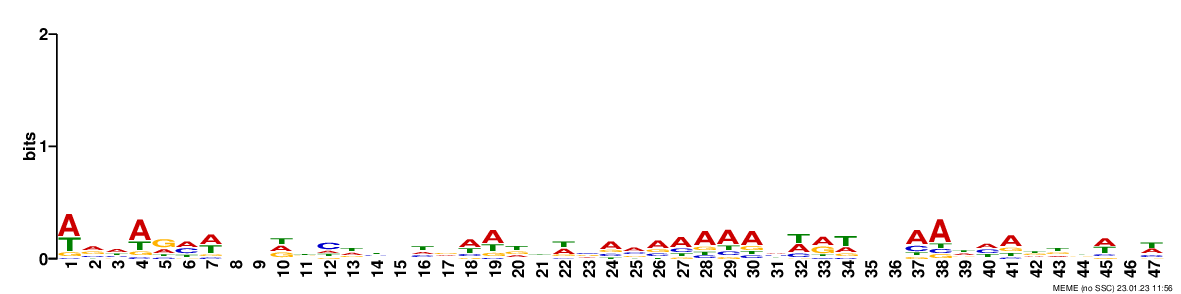

### logo_rc2.png

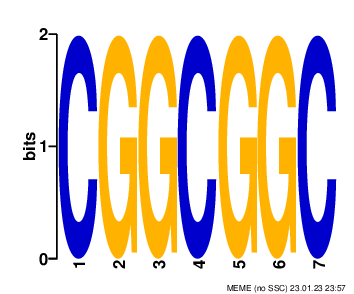

### logo_rc2.png

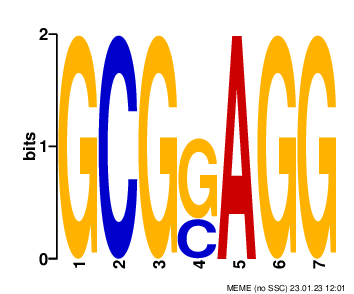

### logo_rc3.png

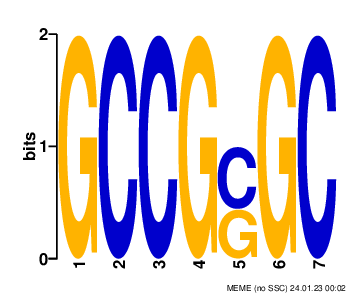

### logo_rc3.png

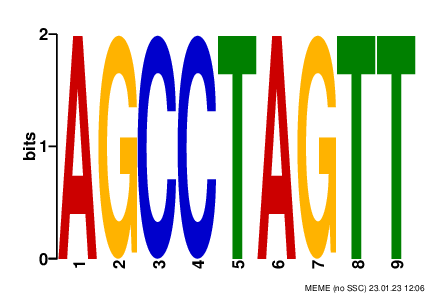

### logo_rc4.png

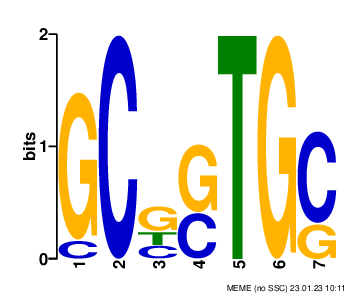

### logo_rc4.png

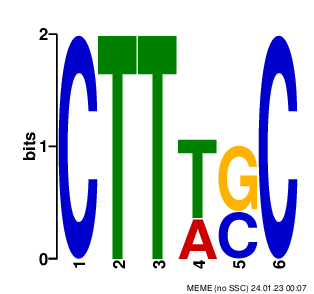

### logo_rc5.png

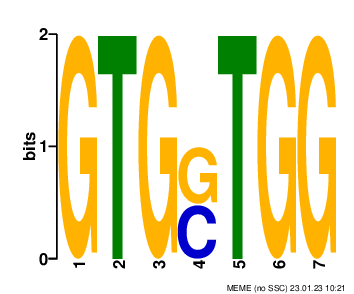

### logo_rc5.png

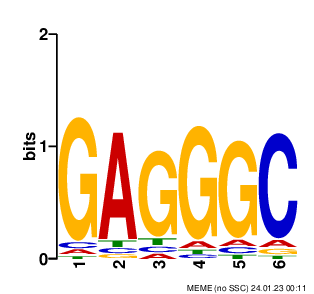

### logo_rc6.png

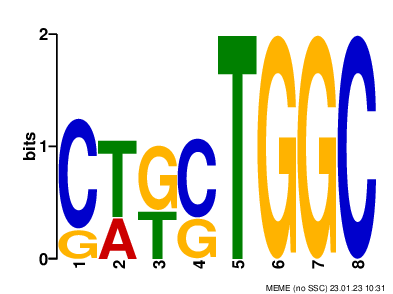

### logo_rc6.png

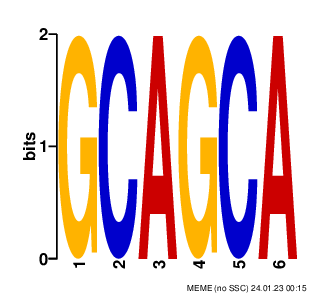

### logo_rc7.png

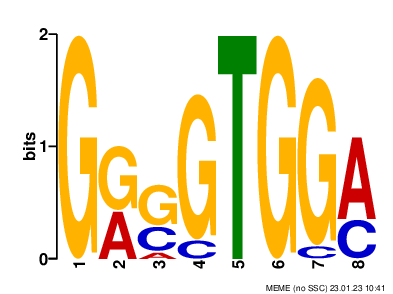

### logo_rc7.png

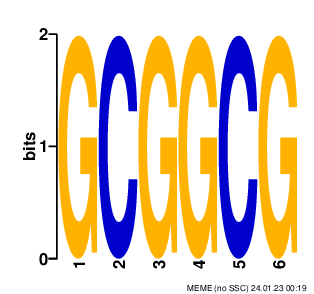

### logo_rc8.png

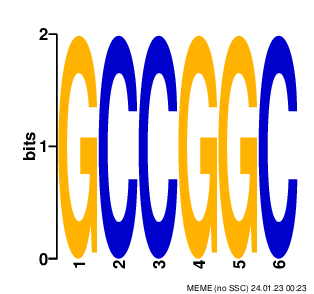

### logo_rc8.png

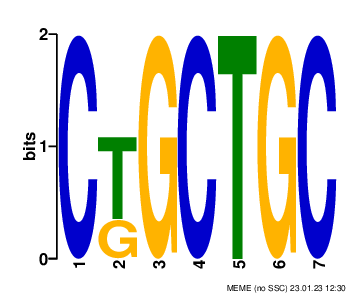

### logo_rc9.png

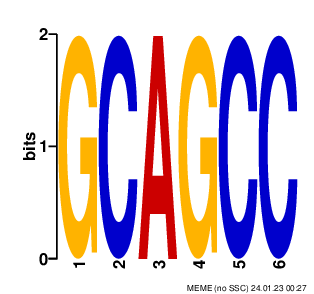

### logo_rc9.png

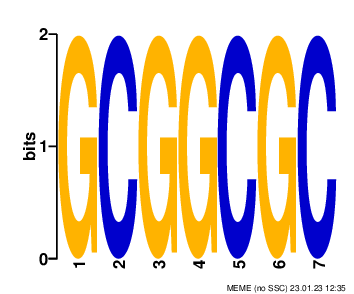

### logo_rc10.png

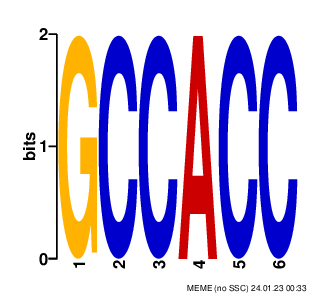

### logo_rc10.png

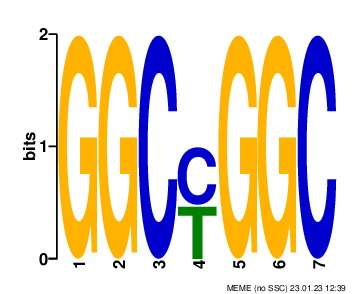

### logo_rc11.png

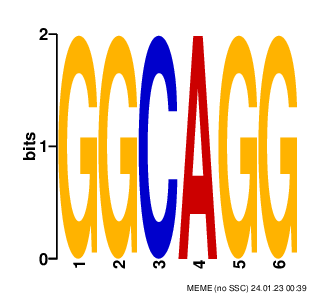

### logo_rc11.png

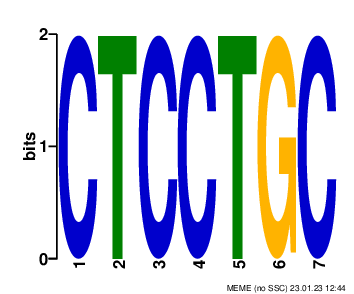

### logo_rc12.png

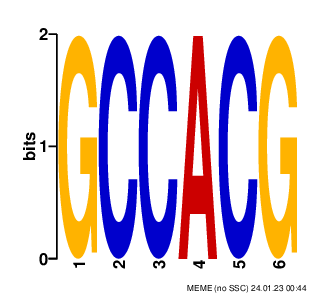

### logo_rc12.png

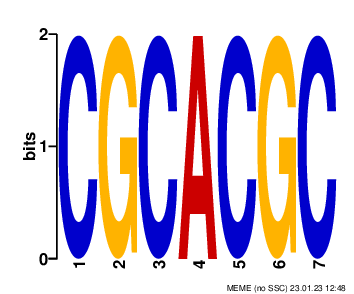

### logo_rc13.png

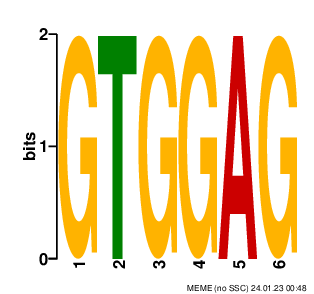

### logo_rc13.png

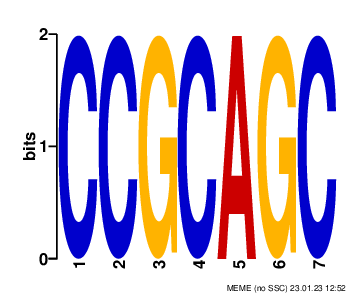

### logo_rc14.png

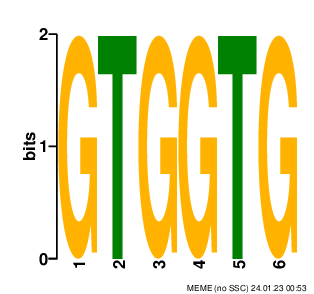

### logo_rc14.png

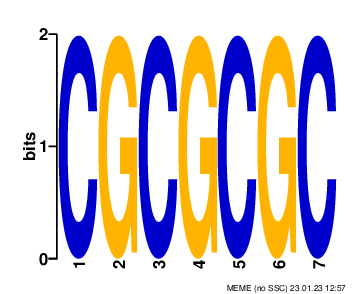

### logo_rc15.png

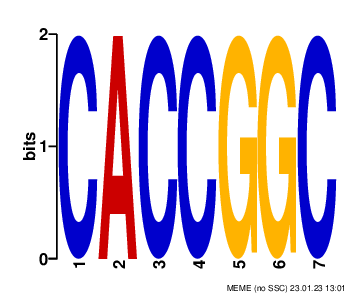

### logo_rc16.png

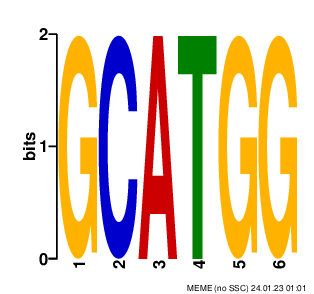

### logo_rc16.png

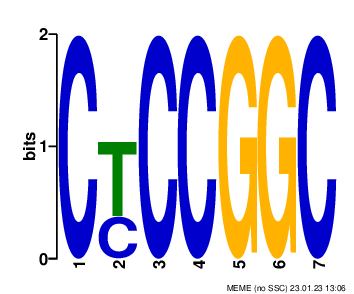

### logo_rc17.png

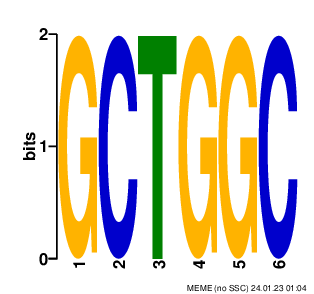

### logo_rc17.png

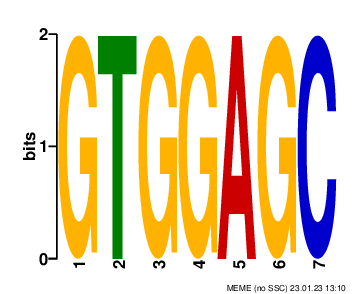

### logo_rc18.png

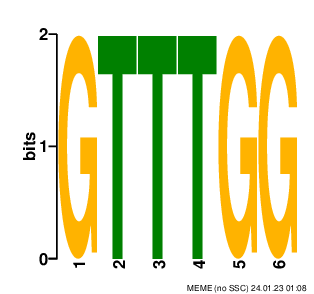

### logo_rc19.png

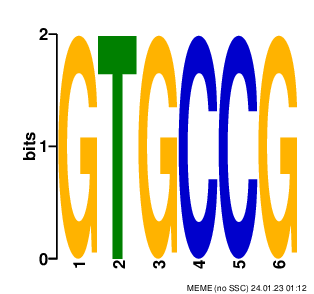

### logo_rc20.png

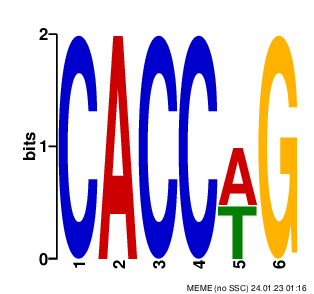

### logo_rc20.png

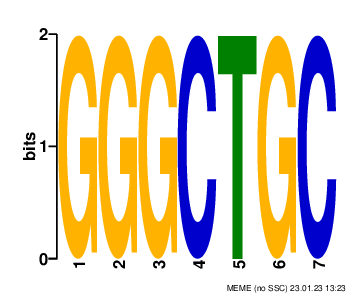

### logo_rc21.png

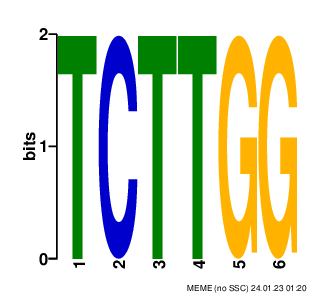

### logo_rc21.png

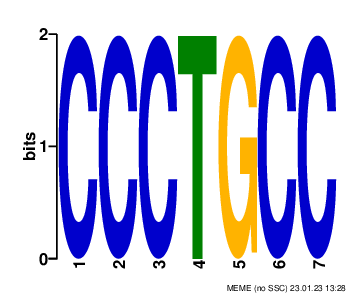

### logo_rc22.png

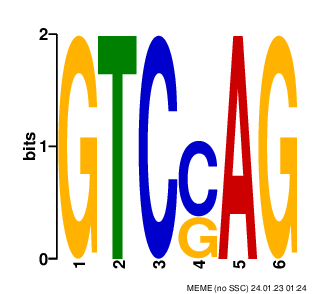

### logo_rc22.png

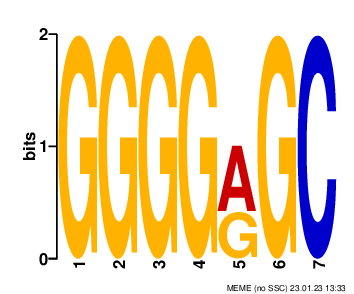

### logo_rc23.png

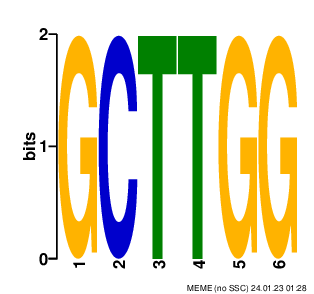

### logo_rc23.png

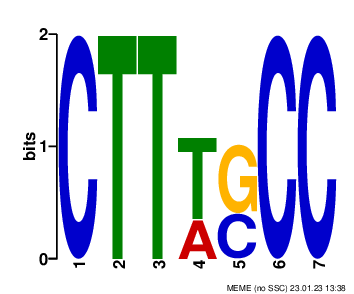

### logo_rc24.png

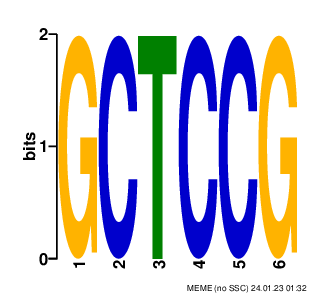

### logo_rc25.png

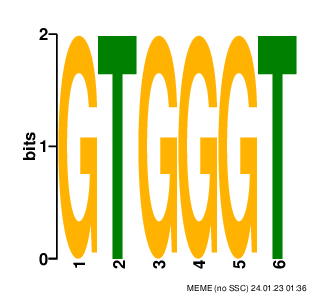

### logo_rc26.png

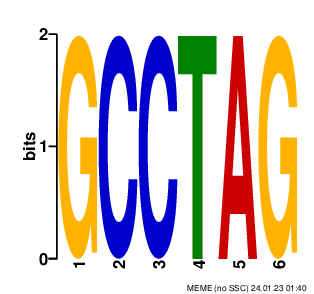

### logo_rc27.png

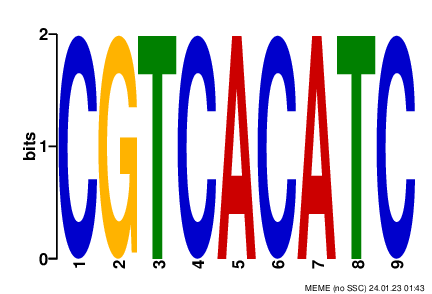

### logo_rc28.png

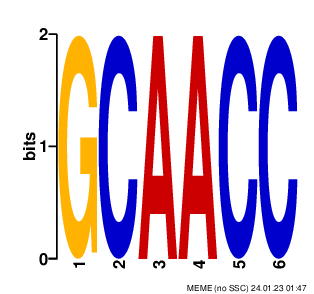

### logo_rc28.png

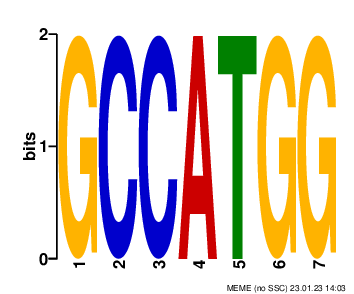

### logo_rc29.png

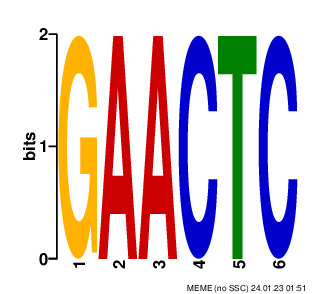

### logo_rc29.png

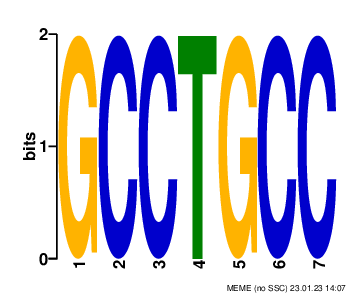

### logo_rc30.png

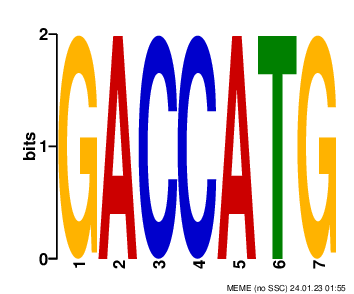
